# Benchmarking cartilage regeneration across ten chondrogenic cell sources using native tissue references

**DOI:** 10.64898/2026.10.05.756631

**Authors:** Simone Ponta, Valerie Pecina, Marielle Airoldi, Daniel Simmen, Eva Novoa Olivares, Thomas Linder, Alexis E. Laurent, Lee Ann Applegate, Gian Salzmann, Valentino F. Bruhin, Martin J. Stoddart, Marcy Zenobi-Wong, Philipp Fisch

## Abstract

A persistent challenge in cartilage tissue engineering is identifying cell sources capable of regenerating cartilage with native-like properties. Variability in chondrogenic potential among human- and animal-derived cells complicates graft development and disease modeling. Here we benchmarked ten chondrogenic cell sources under standardized culture conditions using hyaluronan-alginate biomaterials and implementing a functionality index combining histological and mechanical metrics referenced to native tissue. Mesenchymal stem cells and fetal chondroprogenitors demonstrated superior regenerative capacity, achieving 103% and 95% of native cartilage functionality, respectively. In contrast, adult articular chondrocytes from healthy and osteoarthritic donors produced fibrocartilaginous tissue with functionality scores below 50%. These findings provide a quantitative framework to guide cell source selection and optimize cartilage graft standardization for clinical applications and physiologically relevant disease models, addressing regulatory demands for product characterization.

## Introduction

Cartilage plays a central role in the human body, from load-bearing joints to craniofacial and respiratory structures. Its limited intrinsic capacity for self-repair drives the development of engineered cartilage grafts across a range of clinical applications, including articular cartilage regeneration, auricular and nasal reconstruction, and tracheal repair^1-4^. Among these, joint diseases such as osteoarthritis (OA) represent a particularly large and growing burden. With an aging and increasingly obese population, OA is rising in prevalence and is projected to affect more than one billion people worldwide by 2050^5^, causing chronic pain, reduced mobility, and loss of quality of life, resulting in substantial societal costs^6^. As no disease-modifying drugs are currently available and existing cartilage repair strategies frequently fail to restore long-term tissue function, many OA patients ultimately require total joint replacement^7^. A recurring cause of this failure, and of engineered cartilage graft failure more broadly, is the formation of a fibrocartilaginous phenotype with mechanical properties inferior to native hyaline or elastic cartilage^8^. These limitations have driven the refinement of tissue engineered grafts aiming to recapitulate native physiology for clinical repair and the development of physiologically relevant in vitro models for therapeutic development^9,10^.

A key factor influencing successful cartilage regeneration is the choice of cell source^11,12^, with multiple chondrogenic cell sources available. Their selection is strongly application dependent and their regenerative potential highly variable. Articular chondrocytes are the primary cell type responsible for hyaline cartilage formation and maintenance and have been used clinically through autologous chondrocyte implantation (ACI) and its iterative advancements^13^. However, they often dedifferentiate toward a fibroblast-like phenotype during in vitro monolayer expansion and form a fibrocartilaginous phenotype with mechanical properties inferior to native hyaline cartilage^8,14,15^. Nasal chondrocytes offer a high proliferative capacity, minimal donor-site morbidity, and resilience to inflammatory environments, resulting in the development of nose-to-knee transplantations^16-18^, with clinical outcomes improving when grafts are matured prior to implantation^19^. Auricular chondrocytes provide an accessible elastic-cartilage source with low donor-site morbidity^20-22^, but are limited by the challenge of recapitulating a mature elastic fiber network in vitro. Fetal chondroprogenitor cells have been proposed for off-the-freezer cartilage tissue engineering strategies, with validated chondrogenic attributes in tiered GMP cell banking workflows^23,24^. Infant polydactyly chondrocytes show an enhanced predisposition toward chondrogenic differentiation, but their limited abundance restricts widespread use^25,26^. Mesenchymal stem cells (MSCs) possess multilineage potential and, although their therapeutic effects are often attributed to paracrine signaling and immunomodulation, can undergo robust chondrogenesis^27-31^. Beyond primary human cells, animal-derived chondrocytes, such as bovine, ovine, and porcine, are widely used for their availability and relevance in preclinical models^1,32,33^. Lastly, immortalized chondrocyte-like cell lines such as C28/I2^34^ and ATDC5^35^ offer accessibility and high proliferative capacity but have been reported to show limited chondrogenic capacity^36^.

Despite this diversity and the importance of cell-source selection, functional benchmarks between cell types are rare and often lack tissue-level reference. Studies frequently focus on a single cell source, and comparative analyses remain limited in scope, often restricted to a few sources within a single species or tissue origin (e.g., porcine costal vs. articular chondrocytes, or equine bone-marrow vs. umbilical-cord-blood MSCs)^37,38^. These limitations are particularly relevant in the context of regulatory expectations for tissue engineered products, where consistent characterization and reproducibility across donors are required. In addition, the development of predictive in vitro assays is increasingly emphasized by regulatory agencies including the U.S. Food and Drug Administration (FDA), the European Medicines Agency (EMA), and the UK Medicines and Healthcare products Regulatory Agency (MHRA)^39^. Robust in vitro assessment of engineered cartilage, benchmarked against native tissue, is therefore essential for appropriately evaluating functional graft maturation^12^. We hypothesized that intrinsic differences between chondrocyte sources, such as their developmental origin and differentiation state, determine their capacity to recapitulate the structural and biomechanical properties of native cartilage.

To bridge this gap, we developed a comprehensive functional benchmarking framework for cartilage regeneration to support the development of cartilage grafts for clinical applications and joint-on-chip models for disease modeling. We systematically compare human articular, nasal, auricular and infant polydactyly chondrocytes, fetal chondroprogenitors, mesenchymal stem cells, bovine and ovine chondrocytes, as well as a commonly used chondrogenic cell line (Fig. 1a,d), using standardized culture conditions and biomaterial systems^40^. We benchmarked engineered tissues against native cartilage and established a functionality index to quantify maturation relative to native tissue (Fig. 1b,c)^37^. This study thereby provides a systematic comparison of the chondrogenic capacity across diverse cell sources, enabling identification of cell type-specific performance and revealing cell source-dependent maturation deficits that may guide the development of pharmacological targets to enhance cartilage regeneration.

**Fig. 1.**
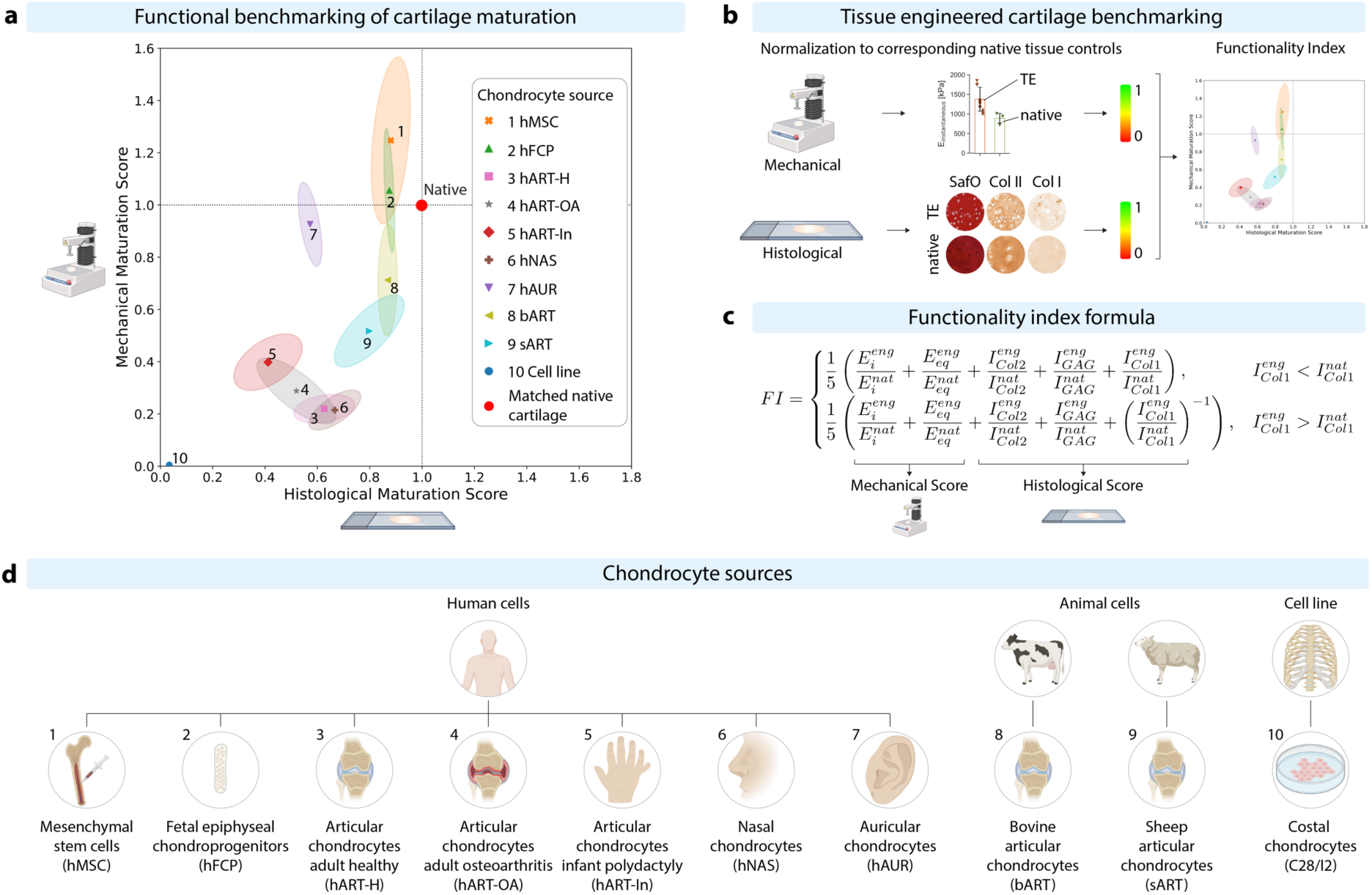
Functionality index and study design. (**a**) Functionality index split into histological and mechanical maturation scores of the ten cell types tested in this study. A value of 1.0 corresponds to the respective native cartilage control which was used to benchmark the engineered cartilage. Details on the control used for benchmarking can be found in the respective results paragraphs. (**b**) Benchmarking of engineered cartilage against native tissue controls. Top: tissue engineered (TE) cartilage samples were tested mechanically and their instantaneous (Ei) and equilibrium (Eeq) modulus benchmarked against native cartilage. Bottom: histological images of tissue engineered cartilage, stained for glycosaminoglycans (SafO), collagen II (Col II) and collagen I (Col I), were color deconvoluted, their staining intensity (I) measured and benchmarked against native cartilage. (**c**) Formula used to calculate the functionality index (FI). Each measurement (Ei, Eeq, IGAG, ICol2, ICol1) of engineered (eng) cartilage was normalized to its native tissue control (nat) and equally weighed in the functionality index. Two formulas were used to account for either more or less collagen I and thereby penalize fibrocartilage formation. (**d**) Overview of the cell types tested in this study together with their species and tissue origin.

## Results

### Mesenchymal stem and progenitor cells generate near-native hyaline cartilage

Among the evaluated cell sources, human bone marrow-derived mesenchymal stem cells (hMSCs) and human fetal epiphyseal chondroprogenitors (hFCP) most closely recapitulated the functional properties of native hyaline cartilage. Combining their biomechanical and histological readouts into the cartilage functionality index yielded scores of 103 ± 15% (hMSC, 3 donors) and 95 ± 9% (hFCP, single donor, ID: FE002) of native hyaline cartilage (Fig. 1a).

hMSCs remained viable (89 ± 7%) and formed grafts with a smooth, cartilage-like morphology after 9 weeks (Fig. S1a-e). Across all three hMSC donors (52 ± 27 years), grafts matured from a partially mature state with a hyaline-like matrix but limited biomechanical function at 3 weeks (instantaneous modulus: E_i_ = 553 ± 181 kPa; equilibrium modulus: E_eq_ = 14 ± 20 kPa) to functionally mature cartilage (Fig. 2a-b). At 9 weeks, the instantaneous (E_i_ = 1378 ± 323 kPa) and equilibrium moduli (E_eq_ = 119 ± 65 kPa) of hMSC-derived cartilage were comparable to healthy human articular cartilage (E_i_ = 895 ± 150 kPa; E_eq_ = 125 ± 69 kPa), corresponding to a mechanical score of 125 ± 32% (Fig. 2c-d). Histologically, grafts showed abundant glycosaminoglycan (GAG) and collagen II deposition with minimal collagen I across all donors, yielding a histological score of 88 ± 8% (Fig. 2e, Fig. S1f-i).

**Fig. 2.**
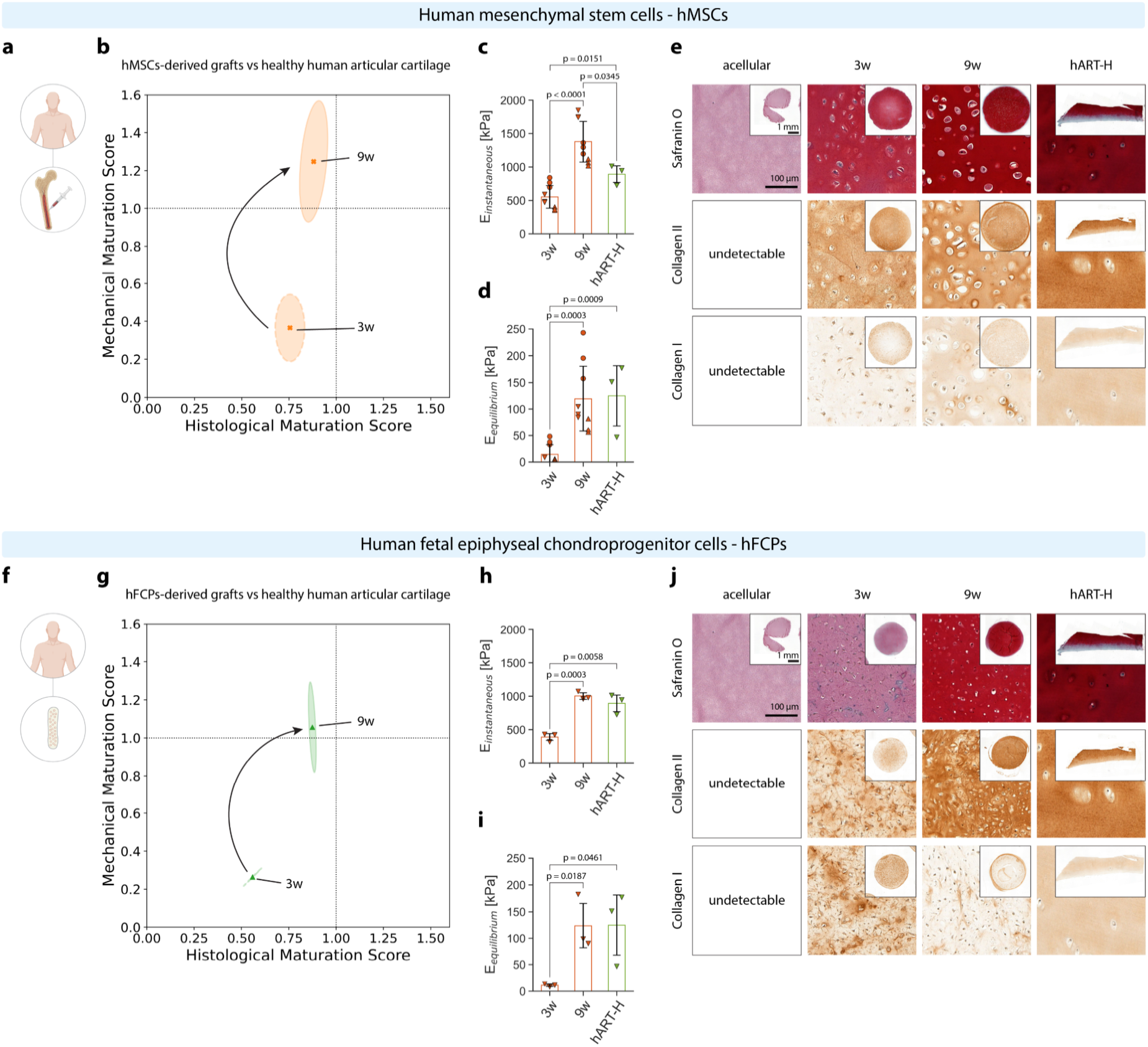
Human mesenchymal stem cells and fetal epiphyseal chondroprogenitors in HATG-Alg. (**a**) Schematic of hMSCs sourced from bone marrow aspirate (52 ± 27 years, 22-73 years, 2 female, 1 male). (**b**) Maturation trajectory of hMSCs from 3 to 9 weeks compared to healthy human articular cartilage. (**c**) Instantaneous and (**d**) equilibrium modulus of grafts at 3 and 9 weeks, compared to native healthy human articular cartilage (hART-H). (**e**) Histological analysis of acellular HATG-Alg control, hMSC grafts matured for 3 and 9 weeks, and healthy human articular cartilage. (**f**) Schematic of hFCPs sourced from the proximal ulnar epiphysis of a 14-week gestation donor. (**g**) Maturation trajectory of hFCPs from 3 to 9 weeks, compared to healthy human articular cartilage. (**h**) Instantaneous and (**i**) equilibrium modulus of grafts at 3 and 9 weeks, compared to native healthy human articular cartilage (hART-H). (**j**) Histological analysis of acellular HATG-Alg control, hFCP grafts matured for 3 and 9 weeks, and healthy human articular cartilage. Scale bar in overview images: 1 mm; scale bar in high-magnification images: 100 µm. hMSC: nd = 3, ns = 3; hFCP: nd = 1, ns = 3.

Similarly, hFCP-derived grafts (Fig. 2f) maintained a high viability (79 ± 4%, Fig. S1j-n) under our designated culture conditions and progressed from a partially mature state (E_i_ = 391 ± 63 kPa; E_eq_ = 11 ± 3 kPa) to functionally mature hyaline cartilage (Fig. 2g). At 9 weeks, grafts reached an instantaneous modulus of 1004 ± 62 kPa and an equilibrium modulus of 124 ± 51 kPa, corresponding to a mechanical score of 106 ± 24% (Fig. 2h-i). Unlike hMSC-derived grafts, hFCP-derived grafts passed through a fibrocartilage-like state at 3 weeks, marked by collagen I deposition and limited GAG accumulation, before maturing into hyaline cartilage (Fig. 2j). By 9 weeks, GAG and collagen II deposition were comparable to native cartilage and collagen I had declined, resulting in a histological score of 88 ± 2% (Fig. S1o-q).

Collectively, hMSC- and hFCP-derived grafts exhibited maturation trajectories towards native hyaline cartilage, underscoring their potential for use in allogeneic cartilage therapies and in vitro models. In both cell types, deposition of extracellular matrix (ECM) components occurred before significant improvements in biomechanical properties, with particularly notable changes in the equilibrium modulus, a measure of the tissue’s ability to resist sustained compression. The observed transition from partially mature to functionally mature cartilage highlights the importance of allowing sufficient time for graft maturation to achieve native-like function. The chondrogenic potential of MSCs is known to vary between donors, with factors such as age, sex, and tissue source contributing to this variability, although findings across studies are inconsistent^41^. In this study, hMSC donors ranged in age from 22 to 73 years and included both sexes, but the sample size was limited to three donors. Larger and more diverse cohorts will be necessary to confirm the generalizability of these near-native functionality scores. hFCPs proliferated more rapidly than hMSCs (Supplementary Results 1, Fig. S2), enabling the creation of large cell banks from a single donor. In contrast, hMSCs generated hyaline-like cartilage across a broad donor age range, providing a wider pool for sourcing cells for allogeneic therapies. The hFCP source used here was derived from a single fetal organ donation (FE002-Cart) that underwent extensive biosafety and functional qualification. Due to their high proliferative capacity and stability, this single hFCP donor could, in principle, be used to manufacture millions of allogeneic cartilage grafts, although further validation at higher cell passages (passages 5-7) in the current culture system is needed. While hFCPs do not require repeated establishment of master cell banks, further functional validation and comparison with donors of different ethnic backgrounds remain important for clinical translation. Overall, the robust expansion, consistent maturation toward native hyaline cartilage, and near-native biomechanical function make hMSCs and hFCPs promising cell sources for cartilage tissue engineering.

### Adult human articular chondrocyte-derived grafts exhibit fibrocartilage-like maturation

Adult human articular chondrocytes are a particularly valuable cell source given their role in joint homeostasis and OA pathophysiology^42,43^. Identifying bottlenecks in their regenerative behavior could inform pharmacological and regenerative treatments for OA. We therefore compared articular chondrocytes from OA patients (hART-OA, 3 donors) with commercially available healthy articular chondrocytes (hART-H, 3 donors). We hypothesized that hART-OA would show reduced regenerative potential owing to senescence and exposure to chronic inflammation. Instead, the two sources converged on similar functionality scores of 43 ± 6% (hART-OA) and 46 ± 8% (hART-H) relative to healthy articular cartilage (Fig. 1a), indicating that the maturation deficit may be intrinsic to the articular lineage rather than driven by OA pathology.

Both, hART-H (58.0 ± 6.6 years; Fig. 3a) and hART-OA (49.0 ± 19.2 years; Fig. 3f) remained viable (hART-H: 95 ± 6%; hART-OA: 84 ± 15%) and retained a homogenous appearance throughout culture (Fig. S3a-e, Fig.S4a-e). The histological score of hART-H was stable throughout culture (3w: 66 ± 11%; 9w: 63 ± 12%; Fig. S4g-i), whereas hART-OA’s score declined significantly (3w: 75 ± 7%; 9w: 52 ± 15%), driven by a loss of GAGs and a rise in collagen I (Fig. S3g-i). Both cell types showed a predisposition toward fibrocartilage, marked by collagen I deposition and limited GAG accumulation (Fig. 3e, j, Fig. S3f, Fig. S4f). Mechanically, both cell types matured substantially, yielding an instantaneous modulus of 415 ± 81 kPa for hART-H (46% of native; Fig. 3c) and 507 ± 185 kPa for hART-OA (57% of native, 895 ± 150 kPa; Fig. 3h), yet their equilibrium modulus remained close to zero (Fig. 3d, i). Since the mechanical score integrates both moduli, this near-zero equilibrium response reduced the overall score, reaching 22 ± 6% (hART-H) and 29 ± 13% (hART-OA) at 9 weeks (Fig. 3b, g). Similar trends were observed when scoring both sources against human OA cartilage (Fig. S5). When expanded with FGF-2+TGF-β3 compared to FGF-2 only, significant shrinkage was observed in both hART-H and hART-OA grafts (Supplementary Results 2, Fig. S6).

**Fig. 3.**
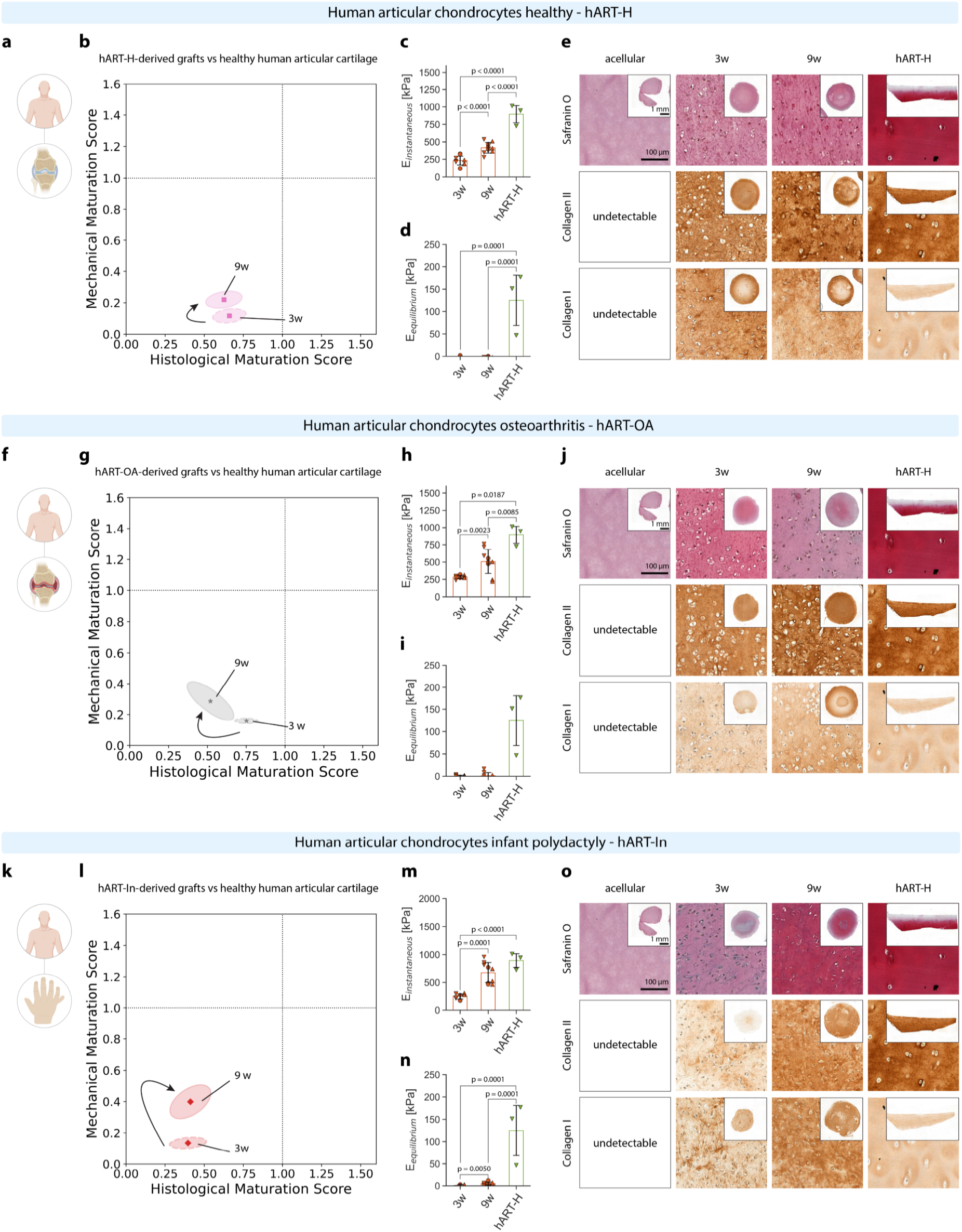
Human articular chondrocytes healthy, osteoarthritis and infant polydactyly chondrocytes in HATG-Alg. (**a**) Schematic of hART-H sourced from healthy knee joints (age: 58.0 ± 6.6 years, age range: 51-64 years, sex: 1 female, 2 males). (**b**) Scoring trajectory of hART-H maturation from 3 to 9 weeks compared to healthy human articular cartilage. (**c**) Instantaneous and (**d**) equilibrium modulus of grafts at 3 and 9 weeks, compared to healthy human articular cartilage (hART-H). (**e**) Histological analysis of acellular HATG-Alg, hART-H grafts matured for 3 and 9 weeks, and healthy human articular cartilage. (**f**) Schematic of hART-OA sourced from osteoarthritic knee joints (age: 49.0 ± 19.2 years, age range: 27-62 years, sex: 2 females, 1 male). (**g**) Scoring trajectory of hART-OA maturation from 3 to 9 weeks compared to healthy human articular cartilage. (**h**) Instantaneous and (**i**) equilibrium modulus of grafts at 3 and 9 weeks, compared to healthy human articular cartilage. (**j**) Histological analysis of acellular HATG-Alg, hART-OA grafts matured for 3 and 9 weeks, and OA human articular cartilage. (**k**) Schematic of hART-In sourced from polydactyly digits (age: 1.3 ± 0.6 years, age range: 1-2 years, sex: 3 males). (**l**) Scoring trajectory of hART-In maturation from 3 to 9 weeks compared to healthy human articular cartilage. (**m**) Instantaneous and (**n**) equilibrium modulus of grafts at 3 and 9 weeks, compared to healthy human articular cartilage. (**o**) Histological analysis of acellular HATG-Alg, hART-In grafts matured for 3 and 9 weeks, and polydactyly human cartilage (hART-In). Scale bar in overview images: 1 mm; scale bar in high-magnification images: 100 µm. hART-H: nd = 3, ns = 3; hART-OA: nd = 3, ns = 3; hART-In: nd = 3, ns = 3.

Collectively, both OA-derived and healthy adult articular chondrocytes converged towards fibrocartilage-like maturation, characterized by collagen I deposition, limited GAG accumulation, and negligible equilibrium mechanics. The emergence of this phenotype despite hMSC- and hFCP-derived grafts achieving near-native cartilage properties under identical culture conditions suggests that the observed maturation deficit may be related to properties intrinsic to articular chondrocytes. Given the central role of proteoglycan-mediated osmotic swelling in cartilage load bearing, insufficient GAG accumulation is a likely bottleneck preventing functional maturation of hART-derived grafts^44^. These findings position hART-derived grafts as a valuable in vitro model to investigate fibrocartilage-associated maturation failure and for developing strategies to restore stable hyaline cartilage regeneration.

### Infant human articular chondrocyte from polydactyly digits form fibrocartilage

While adult articular chondrocytes formed fibrocartilage, we hypothesized that infant articular chondrocytes, harvested from extra digits of infant polydactyly patients (hART-In, 1.3 ± 0.6 years, Fig. 3k), would provide a developmentally younger source of chondrocytes capable of regenerating hyaline cartilage, similar to hFCPs. hART-In are known to express higher levels of hyaline than fibrotic markers and can be passaged longer without losing their chondrogenic phenotype^25,26^. Together with their immunoprivileged status, conferred by the absence of toll-like receptor-4 (TLR4), they represent a promising off-the-freezer source for allogeneic therapies, currently used in Phase III clinical trials^45^. However, when tested under our standardized culture conditions, hART-In formed fibrocartilaginous grafts with a functionality index of 41 ± 11%, comparable to hART-OA and hART-H (Fig. 1a).

hART-In maintained a stable size and remained viable during culture (Fig. S7a-e). Grafts stiffened from 3 to 9 weeks, reaching a mechanical score of 40 ± 11% (Fig. 3l) with an instantaneous modulus of 677 ± 192 kPa at 9 weeks, 76% of that of healthy human articular cartilage (895 ± 150 kPa, Fig. 3m) and significantly higher than the instantaneous modulus of hART-H and hART-OA. However, their equilibrium modulus remained low at 5 ± 3 kPa, corresponding to only 4% of that of healthy human articular cartilage (125 ± 69 kPa, Fig. 3n). Furthermore, grafts showed the deposition of GAGs, collagen II and collagen I at 9 weeks, indicating the formation of fibrocartilage (Fig. 3o, Fig. S7f-i). Unlike hART, hART-In showed a stronger GAG staining, which was however offset by an increased collagen I deposition, yielding only minor changes in the histological score between 3 (40 ± 12%) and 9 weeks (41 ± 13%). Similarly to hART-H and -OA, hART-In were negatively affected by the expansion with FGF-2+TGF-β3, resulting in significant shrinkage and reduced mechanical properties at 9 weeks (Supplementary Results 3, Fig. S8).

In summary, hART-In did not regenerate hyaline cartilage and showed functionality scores similar to those of adult articular chondrocytes. Although grafts derived from hART-In exhibited stronger GAG staining than those from adult articular chondrocytes, this did not result in improved equilibrium mechanical properties. This finding suggests that matrix composition alone is not sufficient to predict functional maturation; factors such as matrix organization likely play a critical role in developing the mechanical properties characteristic of native cartilage. Our results also demonstrate that developmental age by itself does not guarantee successful hyaline cartilage maturation or prevent dedifferentiation toward a fibrocartilage phenotype. The contrast between the regenerative potential of hFCPs and hART-In highlights the need to identify genetic and epigenetic factors that drive these differences in chondrogenic outcomes. However, differences in cell isolation methods (outgrowth culture for hFCPs versus enzymatic digestion for hART-In) and the distinction between prenatal and postnatal origins must be considered when making such comparisons, and dedicated follow-up studies are required. Providing developmentally relevant cues during in vitro maturation may help promote hyaline cartilage formation and reduce the tendency toward fibrocartilage differentiation in chondrocytes.

### Human neural crest-derived chondrocytes produce hyaline and elastic cartilage

Unlike articular chondrocytes, which originate from the mesenchyme, auricular and nasal chondrocytes derive from the cranial neural crest, passing through an ectomesenchyme intermediate before differentiating into chondrocytes^46,47^. Among these, nasal chondrocytes (hNAS) are associated with high proliferation, enhanced chondrogenic capacity and the ability to sustain the local OA inflammatory environment^48,49^. They also offer a less invasive harvest, low donor-site morbidity, and chondrogenic stability. Consequently, nose-to-knee transplantations have gained traction for articular cartilage regeneration and OA treatment^50^. In our study, hNAS reached a functionality index of 48 ± 8% relative to human nasal cartilage (Fig. 1a, 3 donors) and 55 ± 9% relative to healthy human articular cartilage (Fig. S10a-b). While this is comparable to the hART score, hNAS formed hyaline rather than fibrocartilage.

hNAS-derived grafts (24.7 ± 6.1 years; Fig. 4a) maintained a uniform size and high viability throughout culture (Fig. S9a-e). Their instantaneous modulus increased from 240 ± 292 kPa at 3 weeks to 484 ± 150 kPa at 9 weeks (Fig. 4c), corresponding to 43 ± 13% of nasal cartilage (E_i_ = 1134 ± 199 kPa) and 54 ± 17% of healthy articular cartilage (E_i_ = 895 ± 150 kPa). In contrast, negligible maturation of the equilibrium modulus was observed (Fig. 4d), reaching 0.3 ± 2% and 0.5 ± 4% of the equilibrium modulus of nasal and articular cartilage, respectively (Fig. S10b). The lack of equilibrium mechanics thereby affected the overall mechanical score of hNAS-derived grafts, reaching 22 ± 8% (vs. nasal, Fig. 4b) and 27 ± 10% (vs. articular, Fig. S10b). Histologically, grafts transitioned through an initial fibrocartilage state at the outer border of the grafts to a homogenous hyaline cartilage-like state, characterized by GAGs, collagen II and reduced collagen I (Fig. 4e, Fig. S9f-i), resulting in an overall histological score of 67 ± 10% (vs. nasal, Fig. 4b) and 74 ± 11% (vs. articular, Fig. S10b-e). As for the articular sources, hNAS were impaired by the expansion with FGF-2+TGF-β3 (Supplementary Results 4, Fig. S11).

**Fig. 4.**
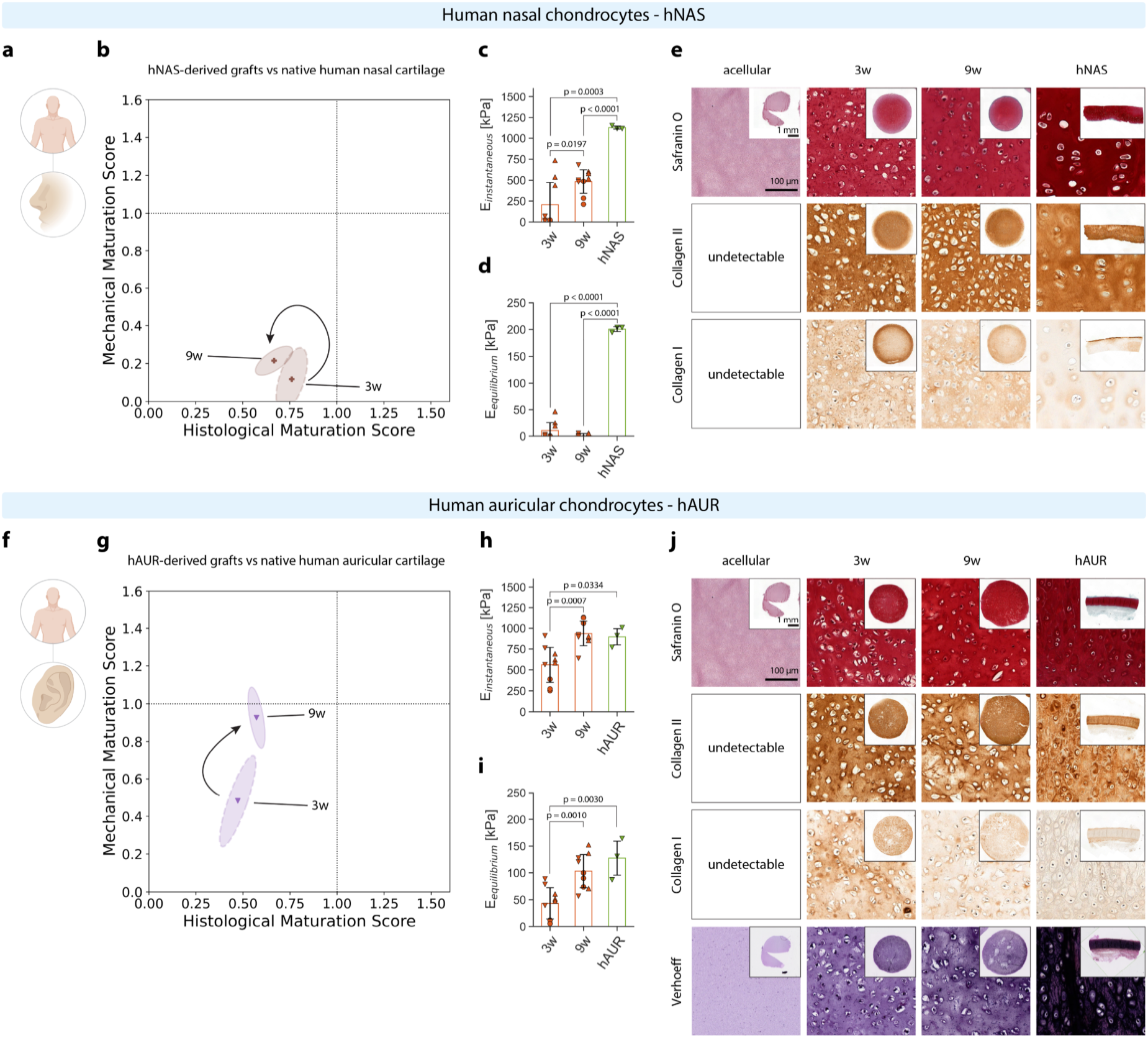
Human nasal and human auricular chondrocytes in HATG-Alg. (**a**) Schematic of hNAS sourced from nasal reconstructive surgery (age: 24.7 ± 6.1 years, age range: 18-30 years). (**b**) Scoring trajectory of hNAS maturation from 3 to 9 weeks compared to human nasal cartilage. (**c**) Instantaneous and (**d**) equilibrium modulus of grafts at 3 and 9 weeks, compared to native human nasal cartilage control (hNAS). (**e**) Histological analysis of acellular HATG-Alg, hNAS grafts matured after 3 and 9 weeks, and human nasal cartilage. (**f**) Schematic of hAUR sourced from otoplasty surgery (age: 11.3 ± 4.7 years, age range: 6-15 years, sex: 2 females, 1 male). (**g**) Scoring trajectory of hAUR maturation from 3 to 9 weeks compared to human auricular cartilage. (**h**) Instantaneous and (**i**) equilibrium modulus of grafts at 3 and 9 weeks, compared to native human auricular cartilage control (hAUR). (**j**) Histological analysis of acellular HATG-Alg, hAUR grafts matured after 3 and 9 weeks, and human auricular cartilage. Scale bar in overview images: 1 mm; scale bar in high-magnification images: 100 µm. hAUR: nd = 3, ns = 3; hNAS: nd = 3, ns = 3.

In contrast to the aforementioned cell sources, auricular chondrocytes (hAUR) form the elastic cartilage of the auricle^51,52^. Their application is therefore specialized to elastic cartilage regeneration, such as microtia, the congenital malformation of the external ear, and reconstructions after burns, cancer or trauma^53,54^. Integrating their biomechanical and histological scores, hAUR converged towards native auricular cartilage with a functionality index of 69 ± 4% compared to human auricular cartilage (Fig.1a, 3 donors).

Like hMSC- and hFCP-derived grafts, hAUR-derived grafts (11.3 ± 4.7 years, Fig. 4f) remained viable and progressively matured towards native tissue properties (Fig. 4g, Fig. S12a-e). At 9 weeks, they reached an instantaneous modulus of 936 ± 155 kPa, and an equilibrium modulus of 103 ± 33 kPa, corresponding to 104% and 80% of human auricular cartilage (E_i_ = 897 ± 119 kPa; E_eq_ = 128 ± 39 kPa, Fig. 4h-i). Histologically, grafts progressed through a fibrocartilaginous state, characterized by collagen I, towards elastic cartilage with a significant increase in elastin and a reduction in collagen I staining intensity (Fig. 4j, Fig. S12f-j). However, although elastin staining intensity increased, the grafts did not reproduce the elastic-fiber organization of native elastic cartilage, reflecting an immature stage of elastogenesis^40,55^. At 9 weeks, hAUR-derived grafts reached a mechanical score of 93 ± 16% and a histological score of 57 ± 5%, which was mainly impacted by the immature elastin network (Fig. 4g).

Although hAUR-derived grafts achieved biomechanical properties comparable to native auricular cartilage, the incomplete formation of an elastin network limited their overall functionality score. Achieving full elastogenesis in tissue-engineered auricular grafts will require a deeper understanding of the genetic programs and matrix changes that drive elastic cartilage formation^55,56^. The biomechanical stability and low donor-site morbidity of auricular chondrocytes make them attractive not only for elastic cartilage applications but also for articular and tracheal cartilage repair^57,58^. In contrast, our functionality index indicates that hNAS-derived grafts require further optimization to achieve functional maturity. Similar to articular chondrocytes, hNAS grafts lacked equilibrium mechanical properties despite displaying near-native histological features, suggesting that insufficient extracellular matrix organization, rather than inadequate matrix deposition, is a limiting factor. Given their distinct developmental origin and transcriptional profile compared to articular chondrocytes, hNAS may benefit from tailored growth-factor supplementation to enhance their extracellular matrix-forming capacity^50^.

### Animal-derived cartilage grafts mature into hyaline-like cartilage

Animal-derived chondrocytes are a readily available, economical cell source with high chondrogenic capacity and fewer ethical constraints, making them particularly relevant for preclinical animal models^12^ and simplified joint-on-chip (JoC) models^10,59^. Both FDA and EMA have issued guidelines for evaluating tissue engineered cartilage grafts in preclinical large-animal studies, such as in sheep, goats, horses, and pigs^31,60-62^. Bovine chondrocytes, in turn, are commonly used in in vitro assays for drug development^63,64^. We therefore evaluated sheep (sART) and bovine (bART) articular chondrocytes as readily accessible cell sources relevant to both preclinical studies and in vitro models. Both cell types confirmed the chondrogenic capacity of animal-derived chondrocytes, reaching functionality indices of 80.6 ± 9% (bART) and 68.6 ± 12.6% (sART) relative to their respective native articular cartilage (Fig. 1a).

bART-derived grafts from approximately 6-month-old calves (Fig. 5a, 3 donors) remained viable and matured into cartilaginous tissue (Fig. 5b, Fig. S13a-e). At 9 weeks, they reached an instantaneous modulus of 703 ± 172 kPa (71 ± 17% of native bovine articular cartilage, 991 ± 96 kPa; Fig. 5c) and an equilibrium modulus of 59 ± 24 kPa (72 ± 29% of native bovine articular cartilage, 83 ± 40 kPa; Fig. 5d), making them one of the few non-MSC/FCP sources to develop substantial equilibrium mechanics, corresponding to a mechanical score of 71 ± 21%. Histologically, grafts closely resembled native tissue across all three donors (Fig. 5e, Fig. S13), with the presence of GAGs, collagen II and absence of collagen I, resulting in a histological score of 87 ± 4%. Hyaline matrix emerged as early as 3 weeks (Ei = 581 ± 235 kPa; Fig. 5c, e), and the equilibrium modulus increased significantly from 18 ± 9 kPa at 3 weeks to 59 ± 24 kPa at 9 weeks (Fig. 5d). Unlike human articular sources (Supplementary Results 2), bART did not lose their regenerative potential when expanded with FGF-2+TGF-β3, reaching 126,184 ± 44,082 cells cm⁻² after 6 days at passage 3 (PDT: 1.13 ± 0.09 days), making bART well suited to high-yield expansion in initial drug screening applications.

**Fig. 5.**
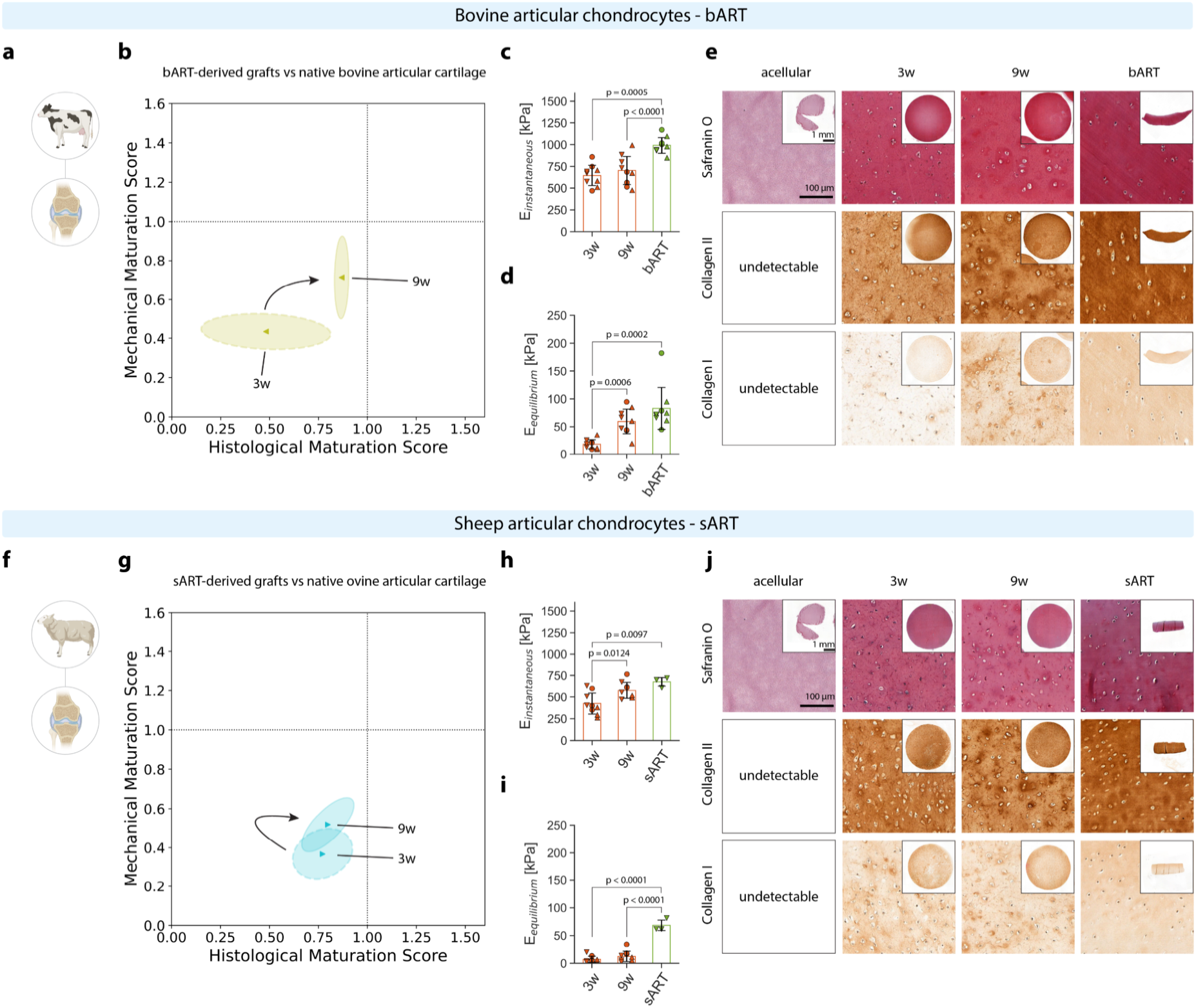
Bovine and ovine chondrocytes in HATG-Alg. (**a**) Schematic of bART sourced from bovine knee joints (∼ 6-month-old calves). (**b**) Scoring trajectory of bART maturation from 3 to 9 weeks compared to bovine articular cartilage. (**c**) Instantaneous and (**d**) equilibrium modulus of the grafts at 3 and 9 weeks, compared to native bovine articular cartilage (bART). (**e**) Histological analysis of acellular HATG-Alg, bART grafts matured for 3 and 9 weeks, and bovine articular cartilage. (**f**) Schematic of sART sourced from sheep knee joints (∼ 3-year-old sheep). (**g**) Scoring trajectory of sART maturation from 3 to 9 weeks compared to ovine articular cartilage. (**h**) Instantaneous and (**i**) equilibrium modulus of the grafts at 3 and 9 weeks, compared to native sheep articular cartilage (sART). (**j**) Histological analysis of acellular HATG-Alg, sART grafts matured for 3 and 9 weeks, and sheep articular cartilage. Scale bar in overview images: 1 mm; scale bar in high-magnification images: 100 µm. bART: nd = 3, ns = 3; sART: nd = 3, ns = 3.

sART-derived grafts (Fig. 5f) followed a different mechanical trajectory (Fig. 5g). Their instantaneous modulus reached 85 ± 15% of the instantaneous modulus of native sheep articular cartilage (engineered: 578 ± 99 kPa; native: 677 ± 60 kPa; Fig. 5h), while their equilibrium modulus remained at only 18 ± 15% of the equilibrium modulus of native tissue (engineered: 12 ± 10; kPa native: 68 ± 11 kPa; Fig. 5i), resembling human articular sources rather than bART. Histologically, sART grafts matured into hyaline cartilage with GAGs, collagen II, and reduced collagen I across all three donors yielding a histological score of 80 ± 14% (Fig. 5j, Fig. S14f-i). During maturation, hyaline cartilage was already present at 3 weeks (histological score: 77 ± 15%), while their mechanical score lagged behind, reaching 37 ± 13% (E_i_= 383 ± 181 kPa, E_eq_ = 7 ± 6 kPa, Fig. 5g). Furthermore, sART reached 54,222 ± 19,693 cells cm^-2^ after 6 days of expansion (PDT:1.71 ± 0.19 days) and remained viable with a uniform size throughout culture (Fig. S14a-e).

In summary, animal-derived articular chondrocytes are an easily accessible cell source capable of regenerating hyaline cartilage in a physiologically relevant manner, suitable for preclinical studies and in vitro models. bART consistently generated native-like cartilage, approaching the mechanical and histological properties of bovine tissue while maintaining rapid expansion. Combined with the availability of bovine joints, these characteristics make bART particularly well suited for high-throughput cartilage engineering, biomaterial screening, and early-stage drug testing before validation in human cells. Future studies are required to determine how closely bovine chondrocytes recapitulate hART inflammatory responses under OA conditions. In contrast, sART reproduced the mechanical maturation profile observed in hART, reaching native-like instantaneous moduli but reduced equilibrium mechanics. Together with the greater similarity of the ovine joint loading environment to humans^65,66^, these findings support the use of sART as a translational model for evaluating regenerative strategies intended for human articular cartilage. Overall, this comparative analysis provides a practical framework for selecting animal-derived chondrocytes according to the intended application, from high-throughput in vitro screening to clinically relevant preclinical studies.

### Immortalized chondrocyte cell line C28/I2 fails to regenerate cartilage

Immortalized chondrocyte-like cell lines are commonly used as in vitro models to study chondrogenesis and cartilage biology^36^. Although they offer the advantage of an extended proliferative capacity and have been employed in metabolic studies, their expression of cell surface markers and ECM components is frequently altered compared to primary chondrocytes^67^. The C28/I2 cell line (single donor) resulted in failed cartilage regeneration with a combined biomechanical and histological functionality index of 2.7 ± 0.7% relative to human hyaline cartilage (Fig. 1a).

While C28/I2 chondrocytes remained viable (Fig. 6a-c), they failed to produce neocartilage. Histological analysis across time points revealed no detectable deposition of glycosaminoglycans (GAGs), collagen II, or collagen I (Fig. 6e), with matrix formation limited to a sparse pericellular collagenous matrix across all samples. In addition to failed ECM production, mechanical testing revealed a compressive modulus of 4 ± 3 kPa (Fig. 6d), corresponding to approximately 0.5% of native human articular cartilage (895 ± 125 kPa).

**Fig. 6.**
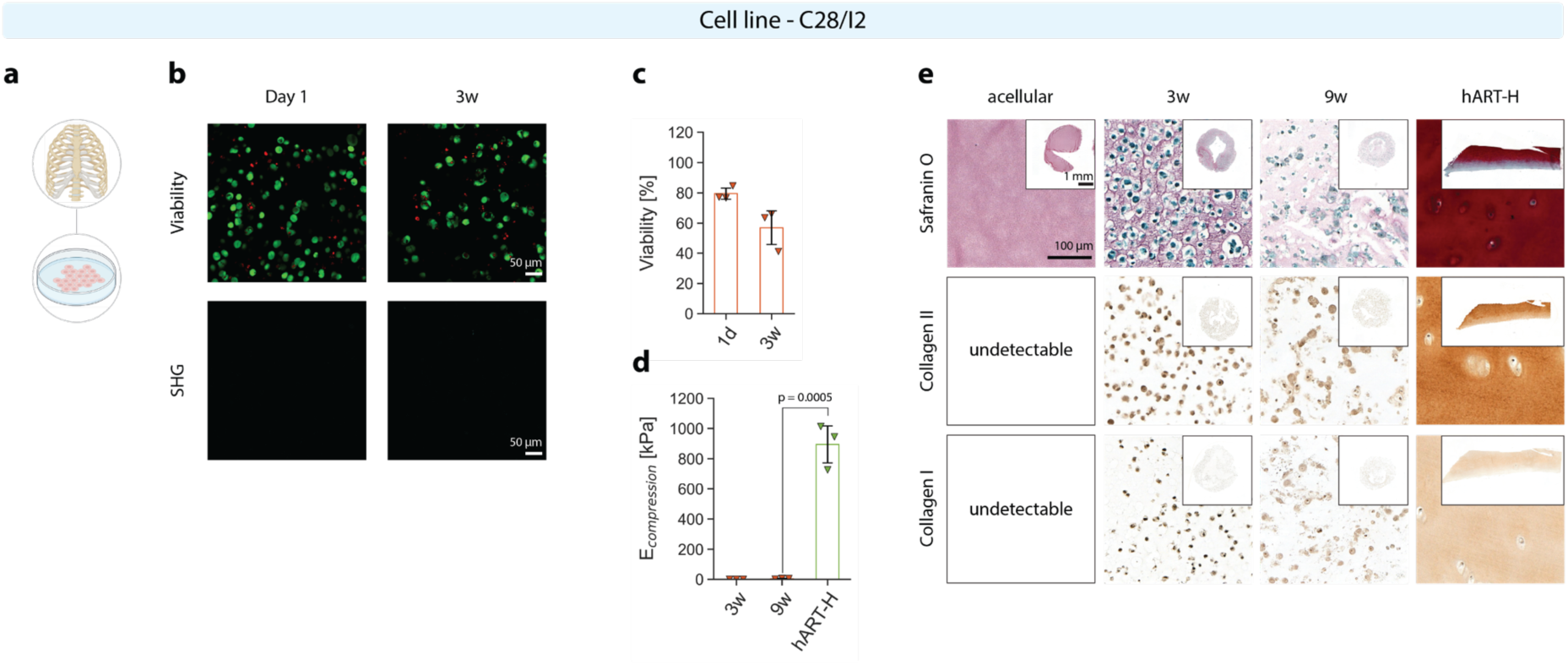
C28/I2 cell line in HATG-Alg. (**a**) Schematic of the C28/I2 cell line, derived from costal cartilage chondrocytes which were subsequently immortalized. (**b-c**) Viability and second harmonic generation of C28/I2 cell line in HATG-Alg at day 1 and 3 weeks. (**d**) Instantaneous modulus of the grafts at 3 and 9 weeks, compared to native healthy human articular cartilage (hART-H). (**e**) Histological analysis of acellular HATG-Alg, C28/I2 grafts matured for 3 and 9 weeks, and healthy human articular cartilage. Scale bar in overview images: 1 mm; scale bar in high-magnification images: 100 µm. nd = 1, ns = 3.

These results indicate a markedly limited capacity of C28/I2 cells to generate functional cartilage under the tested conditions. A possible explanation for the lack of neocartilage formation could be associated with the cell immortalization process itself. The SV40-mediated disruption of p53 and Rb signaling and the resulting transition to S-Phase (proliferation) may impair terminal chondrogenic differentiation and come at the expense of neocartilage synthesis^36,68^. Their limited performance under the tested conditions indicates that C28/I2 cells may be unsuitable for studying chondrogenesis and cartilage regeneration. Considering the low functionality index of C28/I2-derived grafts, our results indicate that alternative chondrocyte sources may represent a more suitable cell source for generating physiologically relevant cartilage models, particularly since protocols to stabilize chondrogenic differentiation during culture and simplified chondrocyte harvesting protocols have been developed^40,69^.

### Material-dependent maturation

We next aimed to make the biomaterial system more accessible to the broader research community by excluding HATG and focusing on alginate-based formulations. To further broaden applicability across biofabrication modalities such as bioprinting, we incorporated hyaluronan (HA) and poly-N-acetyl-D-glucosamine (sNAG) nanofibrils, both previously shown to support chondrogenesis, across a range of material properties^40,70^. We applied the scoring framework developed herein to benchmark the cartilage regeneration potential of hART-H, hART-OA, hNAS, hAUR, bART and sART in three additional alginate-based biomaterials: 1) Alg-HA: 0.25% Alg, 1.5% HA; 2) Alg-sNAG: 0.25% Alg, 2% sNAG; 3) 0.25% Alg, 1.5% HA, 2% sNAG (HATG-Alg: 0.5% HATG, 0.25% Alg). Preliminary experiments on alginate-only materials (0.25% Alg) showed substantial shrinkage of these grafts and were therefore excluded from this study.

Of these materials, HATG-Alg remained the softest material after crosslinking, with a storage modulus (G’) of 2,246 ± 207 Pa and loss modulus (G’’) of 219 ± 17 Pa, followed by Alg-HA (G’ = 3,984 ± 75 Pa, G’’ = 469 ± 18 Pa), Alg-sNAG (G’ = 9,242 ± 985, G’’ = 1,721 ± 166 Pa), and Alg-HA-sNAG (G’ = 11,491 ± 760 Pa, G’’ = 2,370 ± 201 Pa; Fig. 7a). Stress-relaxation tests revealed varying relaxation half-times, with HATG-Alg relaxing fastest (407 ± 264 s), followed by Alg-HA (853 ± 122 s), Alg-HA-sNAG (1,237 ± 50 s) and Alg-sNAG (1,860 ± 696 s; Fig. 7b). Together, these formulations provided a broad range of mechanical environments, spanning from softer viscoelastic hydrogels (HATG-Alg) to mechanically reinforced nanofibril-containing systems (Alg-sNAG and Alg-HA-sNAG). Next, we assessed extrusion printability through shear thinning and shear recovery tests^71,72^. HATG-Alg showed negligible viscosity across shear rates, while Alg-HA displayed zero-shear viscosity behavior, making both unsuitable for bioprinting (Fig. 7c). Alg-sNAG and Alg-HA-sNAG, in contrast, displayed typical shear thinning behavior associated with bioprintable materials, though shear recovery testing revealed that Alg-HA and Alg-sNAG remained too soft in their uncrosslinked state to be printable, while HATG-Alg displayed negligible shear recovery behavior (Fig. 7d). Alg-HA-sNAG, on the other hand, displayed shear recovery behavior similar to our previously published HATG-Alg bioink (0.5% HATG, 0.25% Alg, 1.5% HA, 2% sNAG)^40^. To confirm the rheological results, we 3D bioprinted a knee model (Fig. 7e). Except for Alg-HA-sNAG, none of the materials were able to sustain their shape.

**Fig. 7.**
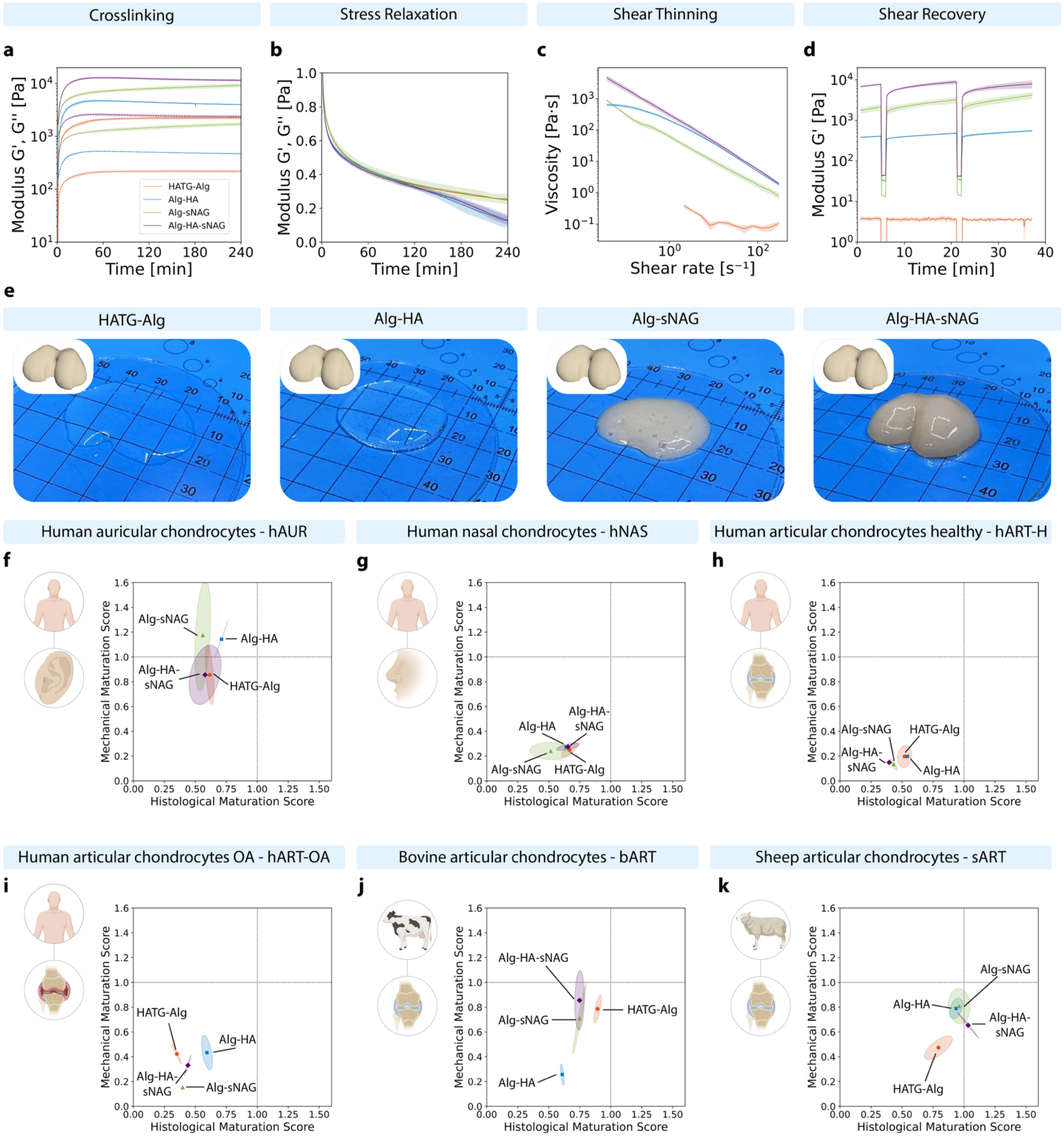
Material characterization and material-dependent maturation of selected cell sources. (**a**) Crosslinking, (**b**) stress relaxation, (**c**) shear thinning and (**d**) shear recovery of HATG-Alg, Alg-HA, Alg-sNAG and Alg-HA-sNAG materials. (**e**) Knee graft bioprinted with the respective material (scale in mm). STL model shown in the upper left corner of each picture. Functionality score at 9 weeks of (**f**) hAUR, (**g**) hNAS, (**h**) hART-H, (**i**) hART-OA, (**j**) bART and (**k**) sART. hAUR, hNAS, bART, sART, hART-H, hART-OA: nd = 1, ns = 3.

We then benchmarked how the different materials affected each cell source and observed distinct responses to the altered biomaterial microenvironment. hAUR-derived grafts showed similar scores across the alternative biomaterials, with Alg-HA resulting in a significantly higher instantaneous modulus, but also significantly smaller graft size compared to HATG-Alg (Fig. 7f, Fig. S16), while hNAS grafts matured similarly across all biomaterials (Fig. 7g, Fig. S17). hART-H showed only minor differences (Fig. 7h, Fig. S18), whereas hART-OA clustered further apart across the different biomaterials, increasing their functionality index from 38 ± 1% in HATG-Alg to 53 ± 5% in Alg-HA, mainly driven by increased GAG deposition (Fig. 7i, Fig. S19). Among the animal-derived cells, no improvements were observed in bART-derived grafts, showing a significant reduction in their score in Alg-HA (Fig. 7j, Fig. S20), whereas sART improved in all biomaterials relative to HATG-Alg (Fig. 7k, Fig. S21). A detailed overview of their mechanical and histological properties can be found in Supplementary Figures S16-21.

In summary, all tested alginate-based biomaterials supported cartilage graft maturation, demonstrating that functional cartilage formation can be achieved using several commercially available materials. This experiment further illustrates how the scoring framework can be used to systematically screen different cell-material combinations and identify opportunities for functional improvement. Notably, the Alg-HA formulation improved the functionality index of hART-OA grafts, while the addition of sNAG nanofibrils enhanced the mechanical properties of grafts derived from hAUR and sART chondrocytes. These results suggest that mechanically reinforcing biomaterials may be particularly advantageous for certain cell sources. Although sNAG nanofibrils were used as a reinforcing additive in this study, their availability is currently limited to a single commercial supplier. Cellulose nanofibrils, which are widely available and well-characterized, have also been used to improve shear-thinning behavior and shape fidelity in alginate-based bioinks for cartilage tissue engineering, and could be substituted to increase the accessibility of this platform^70,73^. While these experiments provide an initial functional assessment of cell-material interactions, further systematic studies are needed to elucidate the molecular mechanisms underlying cell-source-specific responses and to identify optimal biomaterial compositions for each cell population.

## Discussion

In this study, we developed a framework to benchmark the functional regeneration of cartilage and applied it to ten chondrogenic cell sources. Our analysis revealed that hMSCs, hFCPs and hAUR exhibited the highest cartilage regeneration potential, alongside animal-derived bART and sART chondrocytes. hNAS also demonstrated a maturation trajectory towards hyaline cartilage, but their mechanical maturation remained limited. In contrast, hART and hART-In consistently converged towards a fibrocartilaginous phenotype characterized by the lack of GAGs and equilibrium mechanics, while the C28/I2 cell line failed to develop cartilage-like tissue, highlighting substantial differences between immortalized and primary chondrocytes. Taken together, this work provides a broad functional comparison of chondrogenic cell sources with implications for cartilage regeneration and the development of JoC models.

Considering the broad range of cell sources available for cartilage repair, standardized approaches to benchmark the functional quality of engineered grafts are essential. Established approaches rely on histological grading^74-76^ or evaluate total collagen content without discriminating between collagen type I and II^37^. More recently, native tissue-referenced functionality indices have provided a quantitative means of integrating biochemical and biomechanical properties^77^. Proteomics approaches have further demonstrated the ability to resolve temporal changes in the composition of engineered neocartilage and provide a basis for quantitative tissue-quality assessment^33^. The functionality index used here provides a complementary, readily implementable framework that combines biomechanical and histological readouts into a single quantitative metric referenced to native cartilage. By combining instantaneous and equilibrium moduli with the histological assessment of GAGs, collagen I, and collagen II, the score captures both functional properties and tissue phenotype and discriminates between hyaline, elastic and fibrocartilaginous outcomes. In its current form, the functionality index equally weights each of the measurements. Depending on the application area, more weight could be attributed to the mechanical or histological readouts. However, care should be taken to analyze the maturation trajectory and each measurement individually since we observed varying maturation trajectories, e.g. faster mechanical maturation with delayed histological maturation versus histological improvements but lacking mechanical adaptation, depending on the cell type. Its simplicity and reliance on widely accessible assays make it particularly suited for systematic comparisons of diverse cell sources and routine benchmarking of engineered cartilage, while subsequent integration with molecular-level measurements, including proteomics and transcriptomics, enables a more detailed characterization of conditions identified from initial benchmarking.

Replicating native cartilage in vitro remains challenging, as tissue engineered grafts must develop sufficient ECM organization and mechanical properties to withstand the demanding loading environment of synovial joints^78^ and to faithfully replicate cartilage physiology for in vitro studies^79^. In this context, the functionality index introduced herein provides a quantitative benchmark of in vitro tissue quality, distinguishing a fibrocartilaginous phenotype with inferior mechanical properties from hyaline and elastic cartilage. This information becomes important when evaluating at which maturation stage engineered cartilage can be implanted in OA joints, where it needs to withstand the mechanical and inflammatory environment while also efficiently integrating into the surrounding tissue^80,81^. In addition, JoC models to study arthritis and osteoarthritis require capturing the native physiology and pathophysiology of the joint. Cartilage grafts that can faithfully recapitulate native cartilage are therefore needed, together with methods capable of benchmarking them against native healthy and diseased tissue^9^. The functionality index therefore provides a standardized framework for benchmarking engineered cartilage and guiding its optimization, while its predictive value for in vivo regeneration will ultimately require validation in application-specific animal and clinical models.

To achieve functional cartilage regeneration, multiple approaches currently studied in literature can be implemented into the framework presented here. The co-culture of hMSC with hART has been shown to improve the regenerative capacity of hART through paracrine signaling^82^. Together with the cartilage-regeneration potential of hMSC presented here, this may offer a promising strategy to stimulate cartilage regeneration in hART-derived grafts which otherwise mature towards fibrocartilage. In addition to cellular combinations, mechanical loading has been shown to activate mechanosensitive pathways triggering cartilage regeneration^83^. Large scale bioreactors for the stimulation of hART-derived grafts could provide further insights into how mechanical loading stimulates regeneration and ECM arrangement^28^. Upon establishing functional cartilage maturation, these devices would further enable to study how super-physiological loading results in cartilage trauma. Furthermore, while we focused on TGF-β3 for graft maturation throughout this study, studies have highlighted the regenerative potential when combined with other bone morphogenic proteins (BMPs)^84^. These are particularly relevant as developmental studies have shown their importance for cartilage formation during development^85,86^. Lastly, genetic engineering approaches to stimulate targeted protein expression or knock out inhibitory signaling pathways can be incorporated to guide matrix regeneration^26,87^. Overall, the framework presented in this study offers a platform to evaluate cartilage regeneration in mechanistic studies, disease modelling, and therapeutic screenings.

While comprehensive, several limitations exist which should be considered in future studies. The benchmark scoring presented here relies on the semi-quantitative evaluation of histological stainings and is thereby reliant on appropriate native tissue controls. Although efficient to implement for initial screenings of cartilage regeneration, quantitative proteomic and ELISA approaches would offer a valuable tool for inter-study comparability while potentially providing a broader spectrum of information on ECM composition. Beyond ECM-level characterization, transcriptomic and epigenomic approaches could help elucidate the molecular basis of the differences observed across developmental stage, species, and anatomical origin identified in this study, complementing the functional readouts provided here. Furthermore, multiple species and cell types used in recent research, such as chondroprogenitors isolated from adult articular cartilage or induced pluripotent stem cells (iPSCs), were not evaluated in this study. Lastly, the atlas presented here is based on in vitro assessments of cartilage regeneration and does not include in vivo evaluation (i.e., subcutaneous implantation or orthotopic cartilage repair models), which would provide additional insight into graft integration, long-term stability, and host responses.

The work presented here contributes to the standardization of tissue engineered cartilage products and addresses emerging regulatory expectations from agencies including the FDA, EMA, and MHRA. Across these agencies, increasing emphasis is placed on scientifically justified, quantitative approaches for product characterization and on demonstrating product functionality, quality, and consistency. The benchmarking framework provided in this study enables a standardized and simple-to-implement approach to evaluate engineered cartilage relative to native tissue, as well as to compare the regenerative potential between cell sources. The observed differences between cells highlight the distinct regenerative capacities and limitations of individual cell sources and provide a platform to investigate the molecular mechanisms underlying divergent cartilage regeneration outcomes, facilitating the identification of therapeutic targets. Furthermore, by enabling the identification of specific functional deficits relative to native cartilage, the framework provides a rational basis for guiding the optimization of cartilage regeneration strategies, including redirecting maturation away from a fibrocartilaginous phenotype towards native hyaline and elastic cartilage, with the ultimate goal of achieving tissue properties that recapitulate those of native cartilage.

## Acknowledgements

This work was supported by the Swiss National Science Foundation (10000207), Innosuisse (102.321 IP-LS) and the ALIVE initiative (Advanced Engineering with Living Materials) funded by the SFA-AM program (Strategic Focus Area – Advanced Manufacturing) by the ETH Board. The authors gratefully acknowledge ScopeM for their support and assistance in this work. The authors further gratefully acknowledge Marine Polymer Technologies Inc. for providing sNAG nanofibrils and the Tierspital Zürich for providing sheep tissue. We further gratefully acknowledge the S.A.N.T.E and Sandoz Family Foundation for supporting the Swiss progenitor cell transplantation program. Images were created with the support of BioRender (Fisch, P. (2027) https://BioRender.com/zs2idvb).

## Author contributions

- Conceptualization: SP, PF
- Investigation: SP, PF, VP, MA
- Methodology: SP, PF
- Formal analysis: SP, PF
- Resources: DS, ENO, TL, AL, LAA, GS, VFB, MJS
- Supervision: PF, MZW
- Data curation: SP, PF
- Visualization: SP, PF
- Writing – original draft: SP, PF
- Writing – review & editing: all authors
- Funding acquisition: PF, MZW
- Project administration: PF, MZW

## Materials and Methods

### Data Availability

The datasets generated and analyzed in this study are available in the ETH Zurich Research Collection repository under the doi: https://doi.org/10.3929/ethz-c-000806864.

### Cell Source and Ethical Approvals

The C28/I2 human chondrocyte cell line was donated, courtesy of Professor Mary B. Goldring (Hospital for Special Surgery/Weill Medical College of Cornell University, USA)^34^.

Human auricular chondrocytes were isolated from cartilage remnants of otoplasty surgery with three human auricular donors used in this study (age: 11.3 ± 4.7 years, age range: 6-15 years, sex: 2 female, 1 male). Human nasal chondrocytes were isolated form cartilage remnants of nasal reconstruction surgery with three human nasal donors used in this study (age: 24.7 ± 6.1 years, age range: 18-30 years). Informed consent was obtained from the patient or their legal guardian under the ethical license BASEC-Nr.2017–02101.

Healthy human articular chondrocytes were purchased from ACS Biotech (Villeurbanne, France) and three donors were used in this study (age: 58.0 ± 6.6 years, age range: 51-64 years, sex: 1 female, 2 males). Cells from two donors were purchased as isolated cells at passage 0, while the material from the third donor was purchased as a cartilage fragment from which chondrocytes were isolated. The donation, supply and analyzes carried out on biological samples of human origin comply with directives 2004/23/EC (European parliament 2004), 2002/98/EC, 2006/17/EC (European parliament 2006), 2006/86/EC (Parliament 2006).

Human osteoarthritic chondrocytes were isolated from cartilage fragments obtained from joint replacement surgeries of adult patients. Informed consent was obtained from the patients (Kantonale Ethikkommission Zürich, license number PB_2022-01455). Three donors were used in this study (age: 49.0 ± 19.2 years, age range: 27-62 years, sex: 2 females, 1 male).

Human bone marrow aspirates were obtained as surgical waste materials with the patients’ written consent. The Swiss Human Research Act does not apply to research involving anonymized biological material and/or anonymously collected or anonymized health-related data. Therefore, MSCs isolation for this project did not require ethical approval. General consent, which also covers anonymization of health-related data and biological material, was obtained from the patients. Three donors were used in this study (age: 52.3 ± 26.8 years, age range: 22-73 years, sex: 2 females, 1 male).

Primary infant chondrocytes were collected from corrective surgeries of polydactyly patients after informed parents’ consent (Kantonale Ethikkommission Zürich, license number PB_2022-01455). Three donors were used in this study (age: 1.3 ± 0.6 years, age range: 1-2 years, sex: 3 males).

The FE002-Cart primary epiphyseal chondroprogenitor cell source was established from an adapted and modified outgrowth culture from the proximal ulnar epiphysis of a 14-week gestation organ donation, as approved by the Vaud Cantonal Ethics Committee (University Hospital of Lausanne–CHUV, Ethics Committee Protocol #62/07: “Development of fetal cell banks for tissue engineering”, 15. August 2007). The FE002 organ donation was registered under a federal cell transplantation program (i.e., Swiss progenitor cell transplantation program) and the cells used for this study were provided from the privately-owned cell stock of LAA. The epiphyseal cartilage biopsy was collected and processed multimodally for primary cell type establishment and clinical grade therapeutic cell banking^23^. Sufficient parental cell stocks were derived for the subsequent establishment of several hundreds of vials of the master cell bank (MCB) tier of the clinical cell stock, for final use at passage levels 4-7 for optimal chondrogenic responsiveness.

Bovine articular chondrocytes were harvested from knees of ∼ 6-month-old calves, obtained from the local slaughterhouse. Sheep articular chondrocytes were harvested from knees of skeletally mature, ∼ 3-year-old sheep after slaughter, obtained from the Musculoskeletal Research Unit of the University of Zürich. Care was taken to only sample the cartilage from the articular surface.

### Cell Isolation, Culture and Proliferation Metrics

Cells were cultured in a humidified 5% CO_2_ incubator at 37°C. For all experiments, cells were isolated at passage 1 and encapsulated in hydrogels at passage 3, except for the epiphyseal chondroprogenitors, which were encapsulated at passage 4. For passaging, cells were washed once with PBS (Gibco 10010056) and incubated with Trypsin-EDTA (Gibco, 25200056) at 37°C for 5 minutes in the incubator, after which DMEM GlutaMAX™ (Gibco, 41966029) supplemented with 10% FBS (Gibco) was added to block trypsin activity. Cells were transferred to a conical 50 mL Falcon tube and centrifuged at 500 rcf for 5 minutes. Medium was aspirated, and the cell pellet resuspended in fresh expansion medium. Cells were counted and their viability analyzed (Acridine orange/propidium iodide solution, Bucher biotech, F23001) on a LUNA-FX7 Automated cell counter (Logos Biosystems L70001).

Human polydactyly, nasal, auricular, healthy and OA chondrocytes, as well as bovine and sheep articular chondrocytes, were isolated as follows. Cartilage pieces were minced into 1-2 mm pieces, washed in PBS supplemented with antibiotic-antimycotic (Gibco 15240062) and digested overnight in a collagenase solution (DMEM GlutaMAX™, 0.12% w/v collagenase from *Clostridium histolyticum* [Sigma,C9407-100MG], 10% v/v fetal bovine serum [FBS, Gibco, A5256701]) at 37°C under gentle stirring. The cell suspension was subsequently strained through a 100 µm cell strainer and centrifuged at 500 rcf for 5 minutes. The cells were plated at 10,000 cells/cm^2^ and expanded in DMEM GlutaMAX™, 10% v/v FBS, 10 ng/mL fibroblast growth factor 2 (FGF-2, Peprotech, 100-18B) and 10 µg/mL gentamicin (Gibco, 15710049). Auricular cartilage was pre-digested with pronase solution(DMEM GlutaMAX™, 40 U mL^-1^ pronase (pronase from streptomyces, Roche), 2% v/v fetal bovine serum (FBS, Gibco) and 1x antibiotic-antimycotic solution) at 37°C for 90 minutes, prior mincing into 1-2 mm pieces to allow removal of surrounding tissue. hFCPs were originally isolated without antibiotic supplementation and were submitted to trypsin-EDTA only. Cells were expanded in DMEM with added glutamine and 10% FBS until reaching passage level 3 and were subsequently provided in cryopreserved form for this study.

MSCs were isolated via density centrifugation separation and plastic adhesion. The cells were plated at a density of 3,000 cells/cm^2^ and cultured in expansion medium consisting of minimal essential medium without nucleosides (MEMα, Gibco, 22561-054) supplemented with 10% v/v FBS, 10 ng/mL fibroblast growth factor 2 and 10 µg/mL gentamicin.

For subsequent passaging after isolation (i.e., from passage 1 onward), all cell types were expanded in either MEMα (for MSCs) or DMEM GlutaMAX™ (for all remaining cell sources), supplemented with 10% FBS and 10 µg mL^-1^ gentamicin (Gibco). Initially, the expansion medium for all cell types was supplemented with 10 ng mL^-1^ FGF-2 and 10 ng mL^-1^ TGF-β3. However, as this expansion condition negatively affected subsequent chondrogenic differentiation of hMSCs, hART-H, hART-OA, hNAS, and hART-In, these five cell sources were subsequently expanded under two conditions: (i) 10 ng mL^-1^ FGF-2 + 10 ng mL^-1^ TGF-β3 or (ii) 10 ng mL^-1^ FGF-2 alone. Chondrogenic differentiation was then examined following expansion under each condition. Based on these assessments, the expansion condition containing 10 ng mL^-1^ FGF-2 alone was selected for hMSCs, hART-H, hART-OA, hNAS, and hART-In.

For each cell type investigated, we recorded the initial number of cells seeded (N_i_), the final number of cells at harvesting (N_f_) and the time required to reach confluency (Δt). Subsequently, the following cell expansion metrics were calculated: cells/cm^2^ at confluence, averaged expansion time to reach confluency and population doubling time 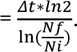 From passage 1 onwards, cells were seeded at 3,000 cells cm^-2^.

### Peptide and Hyaluronan Transglutaminase (HATG) Synthesis

Transglutaminase substrate peptides (glutamine donor peptide: H-NQEQVSPL-ERCG-NH2, lysine donor peptide: Ac-FKGG-ERCG-NH2) were synthesized as reported previously^40^. Briefly, peptides were synthesized on a peptide synthesizer (Prelude X, Gyros Protein Technologies AB) using Rink Amide MBHA resin (RAM-100-HL, Gyros Protein Technologies AB). After synthesis and cleaving, peptides were purified on a preparative HPLC system (Agilent) using an Agilent Prep LC column (C18, 10 µm particle size, 100 Å pore diameter, 250 mm length, 50 mm diameter, AG410910-502, Agilent). Retention peaks from confirmed peptides were used as references and successful synthesis and purification was confirmed by mass spectrometry.

HATG was synthesized as reported previously^21,88^. Briefly, hyaluronan (HA, Lifecore Biomedical) was thiolated with 3,3’-dithiobis(propanoic dihydrazide), vinyl sulfonated with divinyl sulfone and finally functionalized with the glutamine donor peptide (NQEQVSPL-ERCG) or the lysine donor peptide (FKGG-ERCG). The final product was filtered through a 0.2 μm filter and lyophilized. A degree of substitution of ∼10-13% over the disaccharide repeating units was determined by NMR. Rheological crosslinking tests were further carried out and compared to previous synthesis batches to confirm successful synthesis.

### Hydrogel Preparation

All materials used in this study were dissolved in TRIS-buffered glucose (TBG, 200 mM d-glucose, 50 mM TRIS, pH 7.6). To take the cell volume into account, materials were dissolved at a higher concentration to allow the addition of 10% of the final volume in cell volume. A positive displacement pipette was used to aspirate and measure the volume of the cell pellet and the hydrogel-cell mixture was subsequently topped up with TBG to reach final concentrations of: 0.5% w/v HATG, 0.25% w/v Alg, 10 U/mL FXIII (Fibrogammin, CSL Behring), 12.5 U/mL thrombin (TISSEEL, Baxter) for the HATG-Alg gels, 0.25% w/v Alg and 1.5% w/v HA for the Alg-HA gels, 0.25% Alg and 2.0% sNAG nanofibrils (Marine Polymer Technologies, USA) for the Alg-sNAG gels, 0.25% w/v Alg, 1.5% w/v HA and 2.0% w/v sNAG nanofibrils for the Alg-HA-sNAG gels. All materials were tested at a cell concentration of 30 million cells per mL.

### Hydrogel Casting

The hydrogel-cell mixture was cast into sterilized PDMS rings (4 mm diameter, 1 mm height, Sylgard 184, Dow, USA). An atomizer spray bottle was used to spray the casted hydrogel-cell mixture with a 100 mM CaCl_2_ solution to initiate crosslinking. After 15 minutes, 2 mL of 100 mM CaCl_2_ were added to the casted grafts for 45 minutes to finalize crosslinking. After crosslinking, grafts were washed twice in TBG for 5 minutes to remove excess CaCl_2_ and subsequently cultured in chondrogenic medium (DMEM 31966, 50 µg mL^-1^ L-ascorbate-2-phosphate, 40 µg mL^-1^ L-proline, 10 µg mL^-1^ gentamycin, 1% ITS [Insulin-Transferrin-Selenium, Gibco, 41400-045] and 10 ng mL^-1^ TGF-β3). Media changes were performed ever 3-4 days. Grafts were cultured in 12 well-plates with 2 mL of media per graft.

### Bioprinting

Bioprinting was performed on a Bio X bioprinter (Cellink). The 3D model of the human knee was obtained from NIH 3D (Kristen Browne. 2021. 3D Reference Organ for Knee, Male, Right v1.1. https://3d.nih.gov/entries/3DPX-021004. https://doi.org/10.48539/HBM262.NWXL.436). The model was post-processed and scaled in Fusion 360 and sliced on the Bio X bioprinter using a 100% infill pattern. Pneumatic extrusion with a 410 µm nozzle was used for bioprinting. Before printing, the printing pressure was adjusted for each material.

### Rheology

Rheological measurements were carried out on an MCR302e rheometer (AntonPaar) equipped with a Peltier element (P-PTD 200, AntonPaar) and thermal hood (H-PTD 200, AntonPaar). To prevent slipping of the samples, sandpaper was attached to the measurement geometry. Tests were carried out at 25°C and a gap of 500 µm, using a 10 mm diameter parallel plate geometry (PP-10, AntonPaar). A wet cellulose tissue was placed in the thermal hood to prevent sample dehydration.

Crosslinking was tested at 1% shear strain and an angular frequency of 1 rad s^-1^. Data points were collected every 10 s and tests were run for a total of 4 h. Crosslinking was initiated after 1 minute by adding 0.5 mL of 100 mM CaCl_2_ around the sample.

Stress relaxation tests were carried out post-crosslinking. A shear strain of 15% was applied to the sample and held for 6 h, during which the samples were allowed to relax. To determine stress relaxation and the relaxation half-time (τ_½_), the shear stress was normalized to its initial value and the relaxation halftime calculated by calculating the time it took the material to relax to half of the initial stress value minus the final stress value.

Shear recovery tests consisted of 5 intervals. During the 1^st^ interval, a shear strain of 1% at 1 Hz was applied for 5 minutes. Subsequently a high shear interval at 500% shear strain and 1 Hz was applied for 1 minute followed by a recovery interval at 1% shear strain at 1 Hz for 15 minutes. The high shear and recovery interval were repeated once. Data points were collected every 10 s.

During shear thinning tests, the shear rate was logarithmically increased from 0.01 s^-1^ to 300 s^-1^ while data point collection was decreased logarithmically from 300 s per data point to 1 s per data point.

### Compression Testing

Unconfined stress relaxation tests in compression were carried out on a TA.XTplus Texture Analyzer (Stable Micro Systems) equipped with a 500 g load cell. Before each measurement, samples were imaged on a stereomicroscope (Leica Wild Heerbrugg) and the cross-sectional area of the sample was determined in ImageJ2 (v2.16.0/1.54p). Samples were placed between the compression plates and a pre-strain of 15% was applied to ensure full contact between the sample and the plates. Samples were allowed to relax for 10 minutes and 4 consecutive strain intervals were applied. During each interval, the strain was increased by 5% at a strain rate of 5% s^-1^, resulting in a final strain of 20% after the fourth interval. After the first 2 intervals, samples were allowed to relax for 30 minutes and after the last 2 intervals, samples were allowed to relax for 45 minutes. The instantaneous modulus (E_i_) was determined by fitting the stress-strain curve of the last loading interval from 15-20% strain. The equilibrium modulus (E_eq_) was determined by fitting a line through the stress and strain values after relaxation of each interval. The lower limit for reliably measuring the equilibrium modulus was ∼ 10 kPa. Below this value, our analysis method occasionally produced negative equilibrium moduli for extremely soft samples. However, these values were included in our benchmarking approach, as the lack of equilibrium mechanics indicates insufficient extracellular matrix maturation.

### Viability

On day 1 and week 3, 3 grafts from each condition were imaged for viability quantification. The hydrogels were washed 3 times in PBS, stained with 1 µM Calcein-AM (Invitrogen, C3099) and 1 µM propidium iodide (PI, ThermoFisher, P3566) in serum-free DMEM GlutaMAX™ for 30 minutes in a 37°C incubator. The samples were washed 3 times in PBS and imaged using a Leica SP8 Multiphoton microscope with a 25x water immersion objective. To examine total collagen deposition, second harmonic generation (SHG) imaging was performed at 450 nm (detector range: 445-455 nm) with 900 nm two-photon excitation (Mai Tai laser XF, Spectra-Physics).

### Histology and Immunohistochemistry (IHC)

At week 3 and 9, 3 samples per condition were washed in PBS, fixed in 4% paraformaldehyde (PFA, Carl Roth, P087.1) for 2 hours and stepwise dehydrated in a 20%, 40%, 60% and 70% ethanol cascade with 1 hour incubation during each step. Samples were processed and paraffin-embedded using a Milestone Logos J tissue processor (Milestone Medical SRL) and a Paraffin Embedding Station (Medite, TES99). 5 μm sections were cut using a microtome (histocom AG). Before staining, the cut samples were deparaffinized in xylene (VWR, 28975.291) and stepwise rehydrated from ethanol to deionized water. All stained samples were scanned with an automated slide scanner (Panoramic 250, 3D Histech).

#### Safranin O

Slides were first incubated for 5 minutes with Weigert’s Iron Hematoxylin solution (Carl Roth, X906.1 and X907.1), followed by 3 times 30 s wash with deionized water, 2 s in 1% acid alcohol (HCl 37%, VWR 20252.29 diluted in 70% ethanol), 3 times 30 s wash with deionized water, 1 minute 0.02% fast green (Sigma, F7252-5g) and 30 s 1% acetic acid (VWR, 20104.298). The samples were subsequently stained in Safranin O solution (Sigma, S8884-25G) for 30 minutes, washed for 10 s in 95% ethanol, dehydrated to xylene and mounted with Eukitt Quick-hardening mounting medium (Merck, 20104.298).

#### Elastic Stain

Sections were stained according to the manufacturers protocol (HT25, Merck). Briefly, slides were stained in working elastic stain solution for 10 minutes, differentiated in working ferric chloride solutions for 30 s and subsequently dehydrated to xylene and mounted. Between each step, slides were washed in deionized water.

#### Collagens IHC

For colorimetric 3,3′-Diaminobenzidine (DAB) IHC, slides were rehydrated and the samples circled with a hydrophobic pen. Antigen retrieval was performed with 8000 U/mL hyaluronidase (Sigma, H3506-1G) in PBS at 37 °C for 30 minutes. Slides were washed twice in PBS for 5 minutes, and blocked in 5% Bovine Serum Albumin (BSA, Huberlab, A1391,0100) in PBS for 1 hour at ambient temperature. Slides were blotted and incubated with the primary antibody diluted in 1% BSA in PBS overnight at 4 °C. In detail, mouse anti collagen I (1:1000, ab6308, Abcam) and mouse anti collagen II (1:20, II-II6B3, DSHB) antibodies were used. The next day, samples were incubated in 0.3% H_2_O_2_ (Fisher chemicals, H/1750/15) for 15 minutes to block the endogenous peroxidase activity. Slides were then washed twice in PBS for 5 minutes and incubated with a secondary antibody diluted in 1% BSA in PBS (goat-anti-mouse IgG-HRP, 1:1000, ab6789, Abcam) for 1 hour, followed by 3 washes in PBS for 5 minutes. Chromogen development was performed using a DAB substrate kit (Abcam, ab64238) for 5 minutes, followed by 3 washing in deionized water for 30 s. Samples were incubated with Weigert’s Iron Hematoxylin solution for 5 minutes, followed by 3 times 30 s wash with deionized water, 2 s in 1% acid alcohol (HCl 37%, VWR, 20252.29 diluted in 70% ethanol), 3 times 30 s wash with deionized water, 1 minute in bluing agent (0.1% Na_2_CO_3_, Sigma, 451614-25G), 3 times 30 s wash with deionized water. Slides were then dehydrated and mounted with Eukitt Quick-hardening mounting medium.

#### Semi-quantitative evaluation

Semi-quantitative evaluation was performed in ImageJ (Gabriel Landini, https://imagej.net/Colour_Deconvolution, v3.0.2) following our previously developed workflow^70^. Color vectors (Table S1) were generated for Safranin O, Elastic stain, collagen II and collagen I, and images were color-deconvoluted and the mean grey value of the resulting image was measured. After deconvolution, mean grey values were inverted and normalized to native tissue controls. All images of the same staining were processed the same way using an ImageJ macro.

### Functionality Index

The Functionality Index (FI) used throughout this paper combines the mechanical and histological score into one unified measure of engineered cartilage graft maturation (upper index: eng) compared to its native tissue reference (upper index: nat). The following formula was used to calculate the Functionality Index:

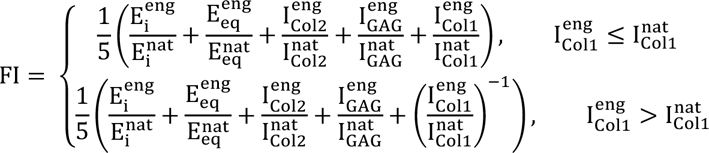

with E_i_ representing the instantaneous modulus, E_eq_ the equilibrium modulus, I_Col2_ the intensity measured from collagen II stainings, I_GAG_ the intensity measured from Safranin O stainings and I_Col1_ the intensity measured from collagen I stainings. The equation was split into two cases in which either more or less collagen 1 was present compared to the native tissue reference to penalize fibrocartilage formation. The respective native tissue reference is mentioned throughout the results section. For elastic cartilage, one additional term based on the elastic staining was included:

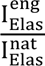

with I_Elas_ representing the intensity measured from Elastic stainings. In addition, the mechanical score (MS) was derived using the following formula:

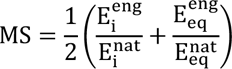

as well as the histological score (HS) using the following formula:

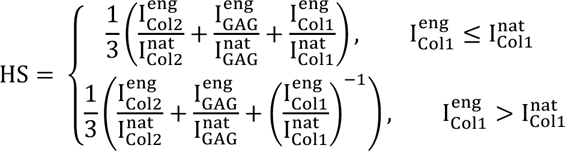

which also included the elastic staining term in the case of human auricular chondrocytes and elastic cartilage.

### Large Language Models

Large language models (ChatGPT, OpenAI and Claude, Anthropic) were used for language editing and proofreading during the preparation of the manuscript in accordance with the guidelines for the responsible use of AI at ETH Zurich (‘Guide for Users of AI at ETH Zurich’, July 24, 2025). No scientific content or analysis was generated using these tools.

### Statistical Analysis

All data are reported as mean ± standard deviation. At each timepoint, grafts were harvested and tested as independent specimens. Statistical analysis was performed in Python (v3.11.5) using the SciPy library (v1.11.1). Independent two-sample t-tests were performed using either the standard Student’s t-test (equal variance) or Welch’s t-test (unequal variance). Levene’s test for homogeneity of variances was used to determine equality of variances. Statistical significance was considered for p-values below 0.05. Unless otherwise stated, each experiment used three biological donors (n_d_ = 3) with three grafts per donor (n_s_ = 3). Fetal epiphyseal chondroprogenitors and the C28/I2 cell line were derived from a single donor (n_d_ = 1, n_s_ = 3), as noted in the corresponding sections. The biomaterial comparison was performed with a single donor per cell source (n_d_ = 1, n_s_ = 3). Shaded ellipses represent ± 1 standard deviation (covariance ellipse).

