## Supplementary Information for "Benchmarking cartilage regeneration across ten chondrogenic cell sources using native tissue references"

#### Supplementary Results

##### 1. Mesenchymal stem and progenitor cells generate near-native hyaline cartilage

Beyond the regenerative capacity, the expansion potential of a cell source influences graft size and the number of in vitro models obtainable from a single donor. To address this, we evaluated the population doubling time as well as the number of cells at harvest at passage 3. hMSCs had a population doubling time (PDT) of 4.8 ± 2.1 days and required approximately 8.7 ± 1.5 days to reach confluency, yielding 12,822 ± 6,140 cells $cm^{-2}$. In contrast, hFCPs proliferated considerably faster, with a 2.8-fold shorter doubling time (PDT = 1.7 days at passage 3) and 50,971 cells $cm^{-2}$ after 7 days of expansion.

Furthermore, not every cell type tolerated expansion with FGF-2+TGF-β3, as opposed to FGF-2 alone. We therefore carried out initial experiments for each cell type that failed to mature under TGF-β3 expansion (see Methods), while keeping the subsequent graft chondrogenic culture phase the same. For hMSCs, TGF-β3 inclusion during expansion impaired subsequent tissue maturation. Although FGF-2+TGF-β3 increased the cell yield (29,714 vs 19,124 cells $cm^{-2}$ for the hMSC donor tested under both conditions), the resulting grafts were significantly softer (Ei = 521 ± 52 kPa; Eeq = 12 ± 5 kPa) than those from hMSCs expanded with FGF-2 alone (Ei = 1284 ± 82 kPa; Eeq = 199 ± 43 kPa; Fig. S2e-f). Consistent with these mechanics, grafts expanded with FGF-2 alone showed higher GAG and lower collagen I content (Fig. S2a-d). In contrast to hMSCs, hFCPs were not negatively impacted by the expansion with FGF-2+TGF-β3, which was consistent with previous reports on the robustness of this cell source under varying culture conditions throughout passage levels 6-7.

Given the established role of TGF-β signaling in fibroblast activation and fibrotic response, combined with the multipotent nature of hMSCs, expansion with TGF-β3 may have driven hMSCs towards a fibrotic rather than a chondrogenic response[89]. In contrast, hFCPs appeared less sensitive to this effect, potentially reflecting their more advanced commitment to the chondrogenic lineage and reduced differentiation plasticity[23].

##### 2. Adult human articular chondrocyte-derived grafts exhibit fibrocartilage-like maturation

Similar to hMSC-derived grafts, hART responded negatively to expansion with FGF-2+TGF-β3, undergoing substantial shrinkage (Fig. S6c-d,i-j). Expanded with FGF-2 alone, hART-H and hART-OA reached 20,940 ± 9,358 cells $cm^{-2}$ (PDT: 2.45 ± 0.63 days) and 15,328 ± 3,237 cells $cm^{-2}$ (PDT: 3.80 ± 1.04 days), respectively. Expansion with FGF-2+TGF-β3 markedly increased expansion rates, with hART-H reaching 92,571 cells $cm^{-2}$ (PDT: 1.21) and hART-OA achieving 69,714 cells $cm^{-2}$ (PDT: 1.54). However, expansion with FGF-2+TGF-β3 also increased collagen I deposition (Fig. S6a-b, g-h). Although biomechanical properties did not differ significantly between the two conditions (Fig. S6e-f, k-l), the pronounced shrinkage after TGF-β3 expansion is a substantial limitation for producing grafts that geometrically match the defect size.

##### 3. Infant human articular chondrocyte from polydactyly digits form fibrocartilage

During expansion with FGF-2, hART-In proliferated significantly faster than hART, reaching 50,804 ± 14,101 cells $cm^{-2}$ after 7 days of culture (PDT: 1.74 ± 0.16 days), a 3.3 and 2.4-fold increase over hART-OA and hART-H, respectively, underscoring their potential as an autologous cell source where large numbers of cells are required. As for hART, expansion with FGF-2+TGF-β3 caused pronounced shrinkage, together with reduced mechanical properties ($E_i$ = 79 ± 59 kPa, $E_{eq}$ = negative value) and lower GAG production (Fig. S8a-f).

##### 4. Human neural crest-derived chondrocytes produce hyaline and elastic cartilage

Although TGF-β3 increased the harvested cell number of hNAS from 43,673 ± 38,728 to 88,210 cells $cm^{-2}$, graft maturation was severely compromised. Histology showed no GAG production over 3 weeks, and the grafts remained too soft for mechanical measurement, so no grafts were collected at 9 weeks (Fig. S11a-f).

hAUR proliferation was comparable to hNAS, reaching 46,457 ± 9,771 cells $cm^{-2}$ at confluency (PDT:1.62 ± 0.35 days). Further expansion conditions were extensively tested in our previous study[40]. The large cell yield makes hAUR well suited to clinical applications such as full-sized human auricular grafts, which could be generated from a 5 mg biopsy.

#### Supplementary Tables

| Staining | Vector | Red | Green | Blue |
|---|---|---|---|---|
| Safranin O | 1 | 0.13271336 | 0.78574693 | 0.6041431 |
| | 2 | 0.2513341 | 0.6987641 | 0.6697462 |
| | 3 | 0.09095116 | 0.63355327 | 0.7683347 |
| Collagen 2 | 1 | 0.16803288 | 0.44223306 | 0.88101923 |
| | 2 | 0.32590333 | 0.53345317 | 0.7805221 |
| | 3 | 0.3267437 | 0.4221379 | 0.8455993 |
| Collagen 1 | 1 | 0.16803288 | 0.44223306 | 0.88101923 |
| | 2 | 0.32590333 | 0.53345317 | 0.7805221 |
| | 3 | 0.3267437 | 0.4221379 | 0.8455993 |
| Elastin | 1 | 0.57578915 | 0.6650111 | 0.47563335 |
| | 2 | 0.49194768 | 0.7324689 | 0.4706132 |
| | 3 | 0.5800034 | 0.6179765 | 0.5307552 |

Table S1 | Color deconvolution vectors used throughout this study to quantify histological images.

### Supplementary Figures

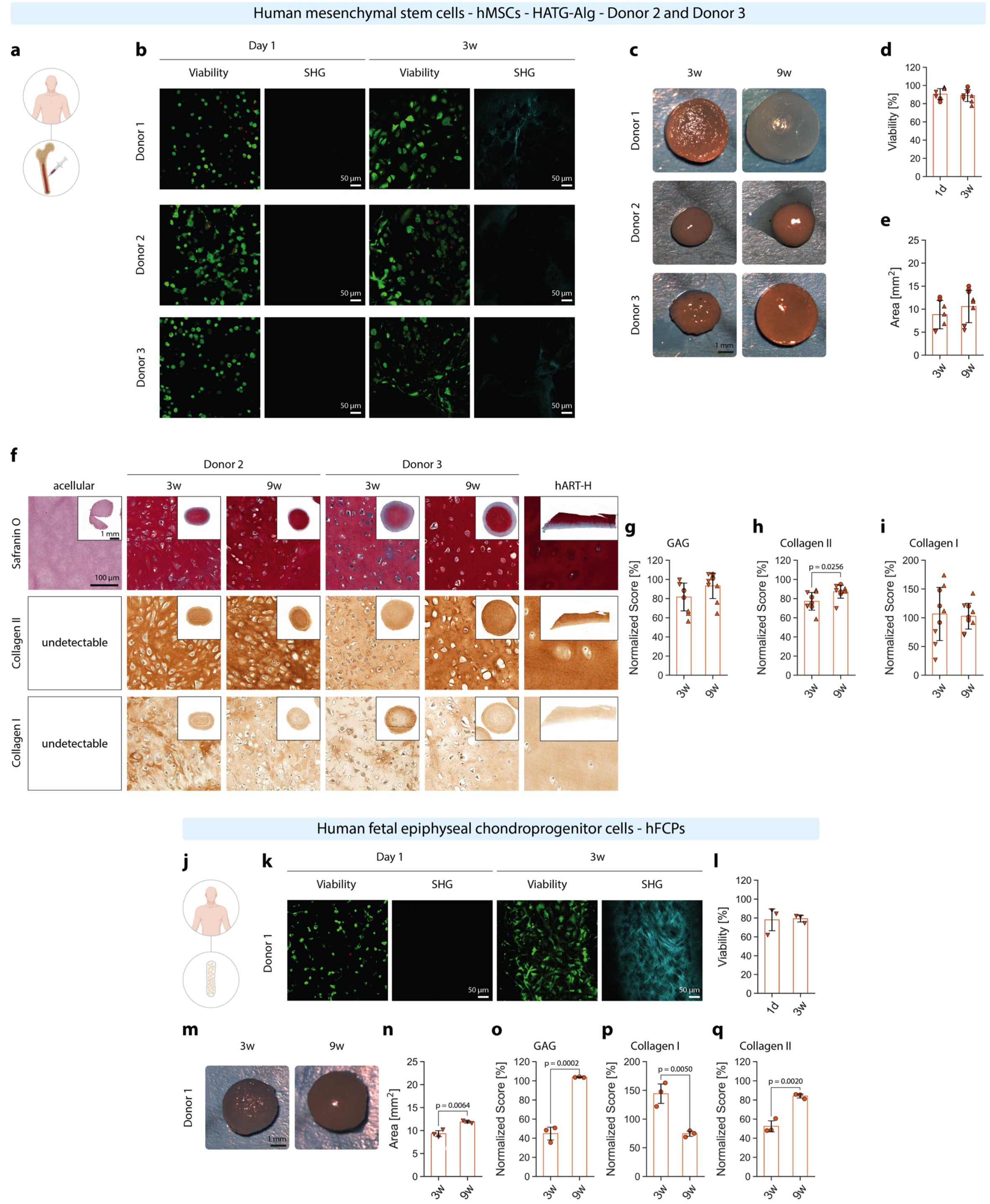


**Fig. S1 | hMSCs and hFCPs in HATG-Alg.** (**a**) Schematic of hMSCs sourced from bone marrow aspirate (52 ± 27 years, min: 22 years, max: 73 years, 2 females, 1 male). (**b**) Viability and second harmonic generation of hMSC grafts at day 1 and 3 weeks and (**d**) viability quantification. (**c**) Gross appearance of hMSC grafts after 3 and 9 weeks and (**e**) graft area. (**f**) Histological analysis of acellular HATG-Alg, MSC grafts matured after 3 and 9 weeks, and healthy human articular cartilage (hART-H). (**g**) GAG, (**h**) collagen II and (**i**) collagen I histological score breakdown between 3- and 9-weeks samples. (**j**) Schematic of hFCPs sourced from the proximal ulnar epiphysis of a 14-week-gestation organ donation. (**k**) Viability and second harmonic generation of hFCP grafts at day 1 and 3 weeks and (**l**) viability quantification. (**m**) Gross appearance of hFCP grafts after 3 and 9 weeks and (**n**) graft area. (**o**) GAG, (**p**) collagen II and (**q**) collagen I histological score breakdown between 3- and 9-weeks samples. Scale bar in viability and second harmonic generation images: 50 µm; scale bar in gross pictures of the samples and in histological overview images: 1 mm; scale bar in high-magnification images: 100 µm. hMSC: $n_d$ = 3, $n_s$ = 3; hFCP: $n_d$ = 1, $n_s$ = 3.

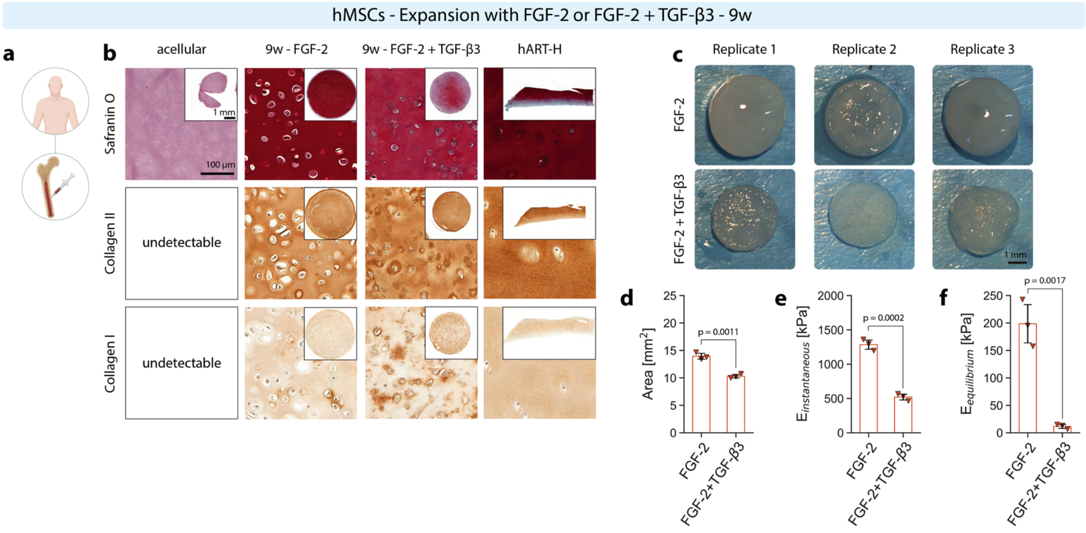


**Fig. S2 | hMSCs expanded with FGF-2 and with FGF-2+TGF-β3 in HATG-Alg.** (**a**) Schematic of hMSCs sourced from bone marrow aspirate (52 ± 27 years, min: 22 years, max: 73 years, 2 females, 1 male). (**b**) Histological analysis of hMSC grafts matured for 9 weeks after expansion with FGF-2 (10 ng mL$^{-1}$) or FGF-2+TGF-β3 (both 10 ng mL$^{-1}$), compared to acellular HATG-Alg and healthy human articular cartilage (hART-H). (**c**) Gross appearance of hMSC grafts after expansion with FGF-2 or FGF-2+TGF-β3 and (**d**) graft area. (**e**) Instantaneous and (**f**) equilibrium moduli of hMSC grafts after expansion with FGF-2 or FGF-2+TGF-β3. Scale bar in gross pictures of the samples and in histological overview images: 1 mm; scale bar in high-magnification images: 100 µm. $n_d = 1$, $n_s = 3$.

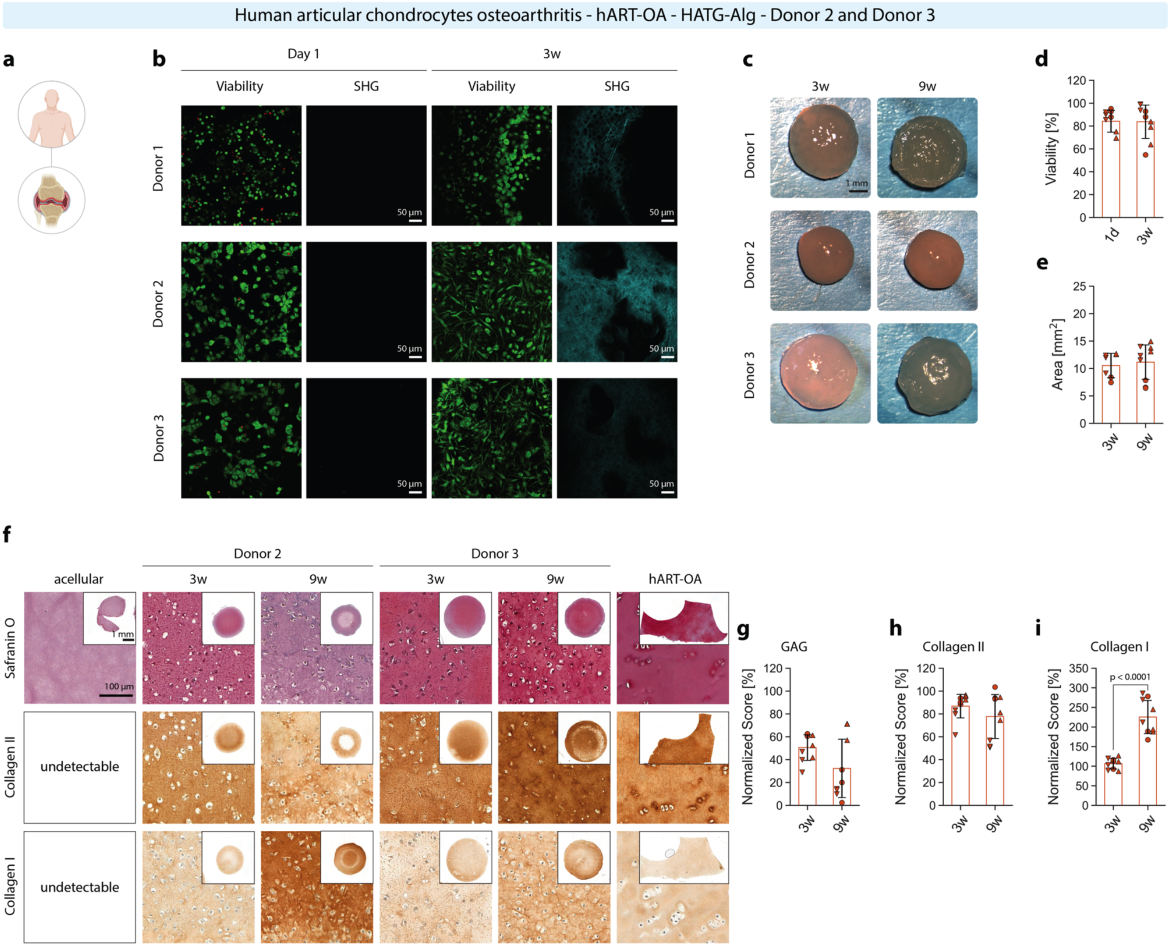


**Fig. S3 | hART-OA in HATG-Alg.** (**a**) Schematic of hART-OA sourced from osteoarthritic knee joints (age: 49.0 ± 19.2 years, age range: 27-62 years, sex: 2 females, 1 male). (**b**) Viability and second harmonic generation of hART-OA grafts at day 1 and 3 weeks and (**d**) viability quantification. (**c**) Gross appearance of hART-OA grafts after 3 and 9 weeks and (**e**) graft area. (**f**) Histological analysis of acellular HATG-Alg, hART-OA grafts matured after 3 and 9 weeks, and OA human articular cartilage (hART-OA). (**g**) GAG, (**h**) collagen II and (**i**) collagen I histological score breakdown between 3- and 9-weeks samples. Scale bar in viability and second harmonic generation images: 50 µm; scale bar in gross pictures of the samples and in histological overview images: 1 mm; scale bar in high-magnification images: 100 µm. hART-OA: $n_d$ = 3, $n_s$ = 3.

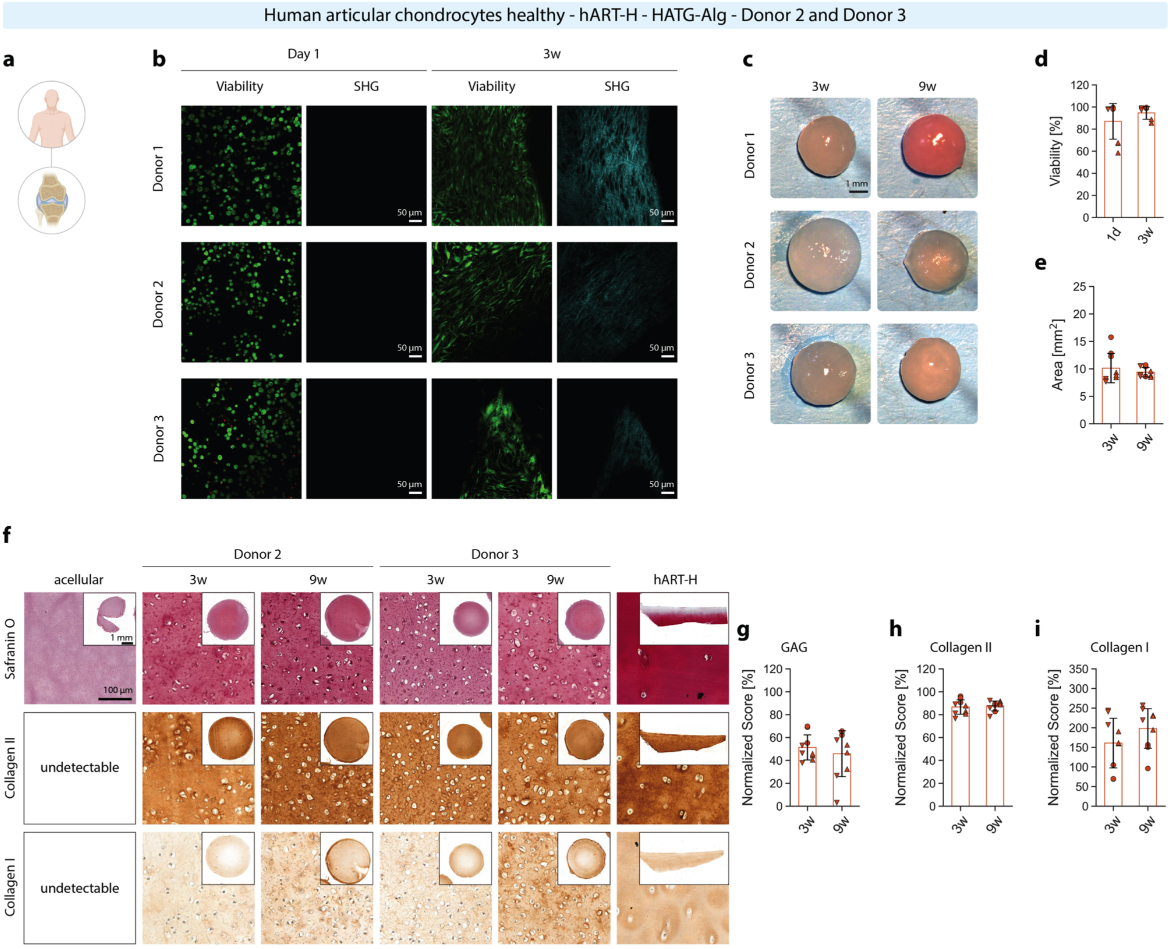


**Fig. S4 | hART-H in HATG-Alg.** (**a**) Schematic of hART-H sourced from healthy knee joints (age: 58.0 ± 6.6 years, age range: 51-64 years, sex: 1 female, 2 males). (**b**) Viability and second harmonic generation of hART-H grafts at day 1 and 3 weeks and (**d**) viability quantification. (**c**) Gross appearance of hART-H grafts after 3 and 9 weeks and (**e**) graft area. (**f**) Histological analysis of acellular HATG-Alg, hART-H grafts matured after 3 and 9 weeks, and healthy human articular cartilage (hART-H). (**g**) GAG, (**h**) collagen II and (**i**) collagen I histological score breakdown between 3- and 9-weeks samples. Scale bar in viability and second harmonic generation images: 50 µm; scale bar in gross pictures of the samples and in histological overview images: 1 mm; scale bar in high-magnification images: 100 µm. hART-H: $n_d$ = 3, $n_s$ = 3.

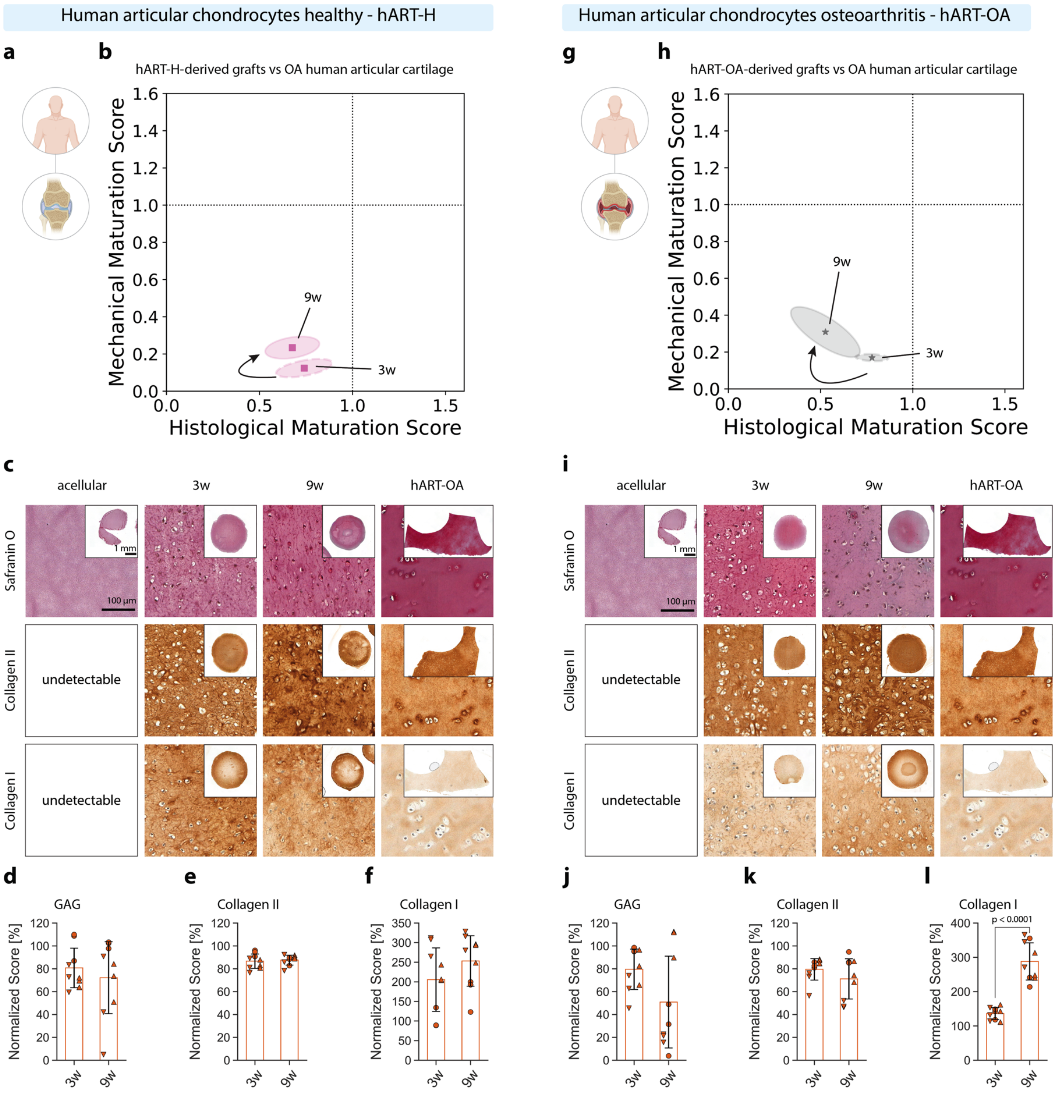


**Fig. S5 | hART-H and hART-OA – Maturation trajectory vs native human OA cartilage.** (**a**) Schematic of hART-H sourced from healthy knee joints (age: 58.0 ± 6.6 years, age range: 51-64 years, sex: 1 female, 2 males). (**b**) Scoring trajectory of hART-H maturation from 3 to 9 weeks compared to OA human articular cartilage. (**c**) Histological analysis of acellular HATG-Alg, hART-H grafts matured after 3 and 9 weeks, and OA human articular cartilage (hART-OA). (**d**) GAG, (**e**) collagen II and (**f**) collagen I histological score breakdown between 3- and 9-weeks hART-H samples normalized to OA human articular cartilage. (**g**) Schematic of hART-OA sourced from osteoarthritic knee joints (age: 49.0 ± 19.2 years, age range: 27-62 years, sex: 2 females, 1 male). (**h**) Scoring trajectory of hART-OA maturation from 3 to 9 weeks compared to OA human articular cartilage. (**i**) Histological analysis of acellular HATG-Alg, hART-OA grafts matured after 3 and 9 weeks, and OA human articular cartilage. (**j**) GAG, (**k**) collagen II and (**l**) collagen I histological score breakdown between 3- and 9-weeks hART-OA samples normalized to OA human articular cartilage. Scale bar in overview images: 1 mm; scale bar in high-magnification images: 100 µm. hART-H: $n_d = 3$, $n_s = 3$; hART-OA: $n_d = 3$, $n_s = 3$.

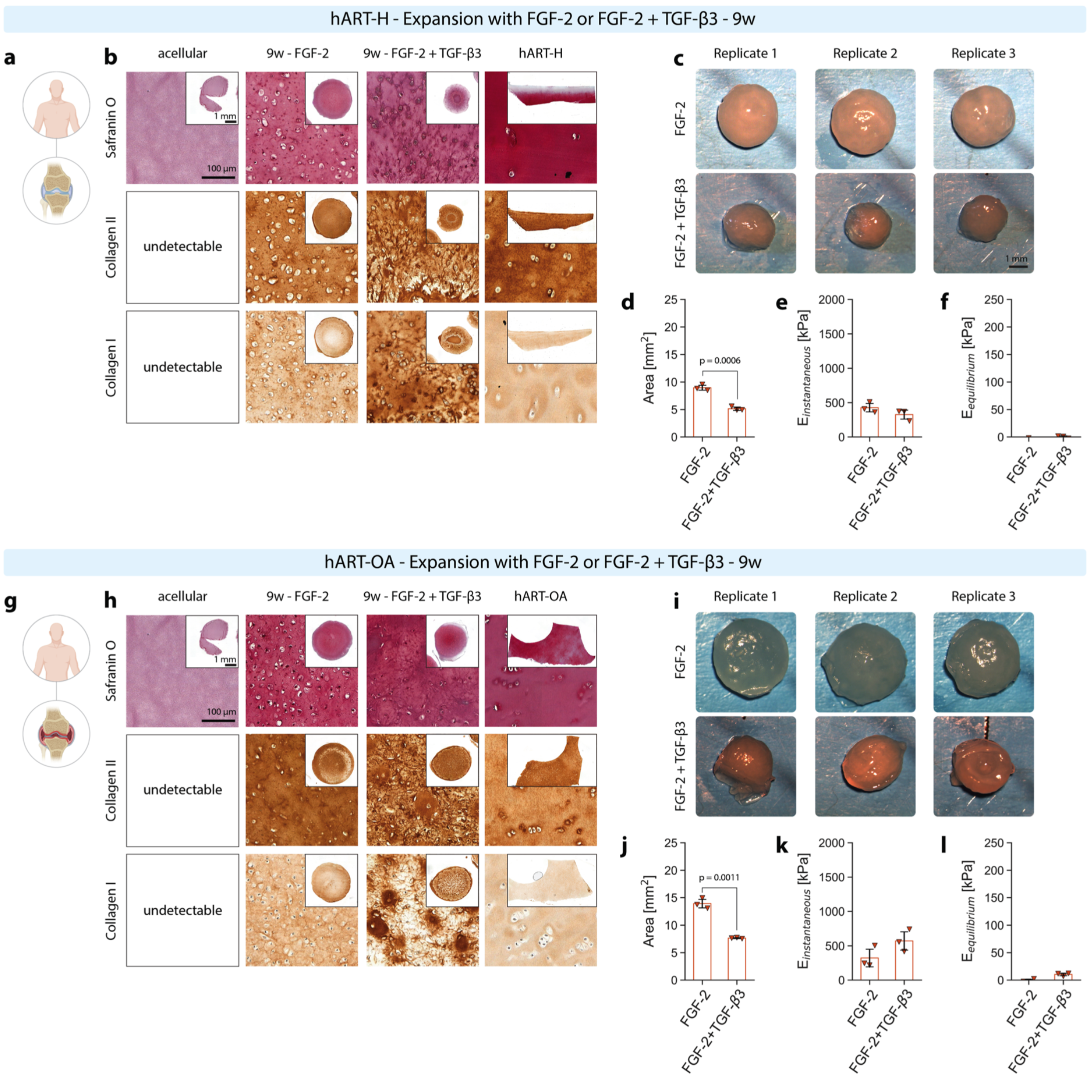


**Fig. S6 | hART-H and hART-OA expanded with FGF-2 and with FGF-2+TGF-β3 in HATG-Alg.** (**a**) Schematic of hART-H sourced from healthy knee joints (age: 58.0 ± 6.6 years, age range: 51-64 years, sex: 1 female, 2 males). (**b**) Histological analysis of hART-H grafts matured for 9 weeks after expansion with FGF-2 10 ng mL$^{-1}$ or FGF-2+TGF-β3 (both 10 ng mL$^{-1}$), compared to acellular HATG-Alg and healthy human articular cartilage (hART-H). (**c**) Gross appearance of hART-H grafts after expansion with FGF-2 or FGF-2+TGF-β3 and (**d**) graft area. (**e**) Instantaneous and (**f**) equilibrium moduli of hART-H grafts after expansion with FGF-2 or FGF-2+TGF-β3. (**g**) Schematic of hART-OA sourced from osteoarthritic knee joints (age: 49.0 ± 19.2 years, age range: 27-62 years, sex: 2 female, 1 male). (**h**) Histological analysis of hART-OA grafts matured for 9 weeks after expansion with FGF-2 (10 ng mL$^{-1}$) or FGF-2+TGF-β3 (both 10 ng mL$^{-1}$), compared to acellular HATG-Alg and human OA articular cartilage (hART-OA). (**i**) Gross appearance of hART-OA grafts after expansion with FGF-2 or FGF-2+TGF-β3 and (**j**) graft area. (**k**) Instantaneous and (**l**) equilibrium moduli of hART-OA grafts after expansion with FGF-2 or FGF-2+TGF-β3. Scale bar in gross pictures of the samples and in histological overview images: 1 mm; scale bar in high-magnification images: 100 µm. hART-H: $n_d = 1$, $n_s = 3$; hART-OA: $n_d = 1$, $n_s = 3$.

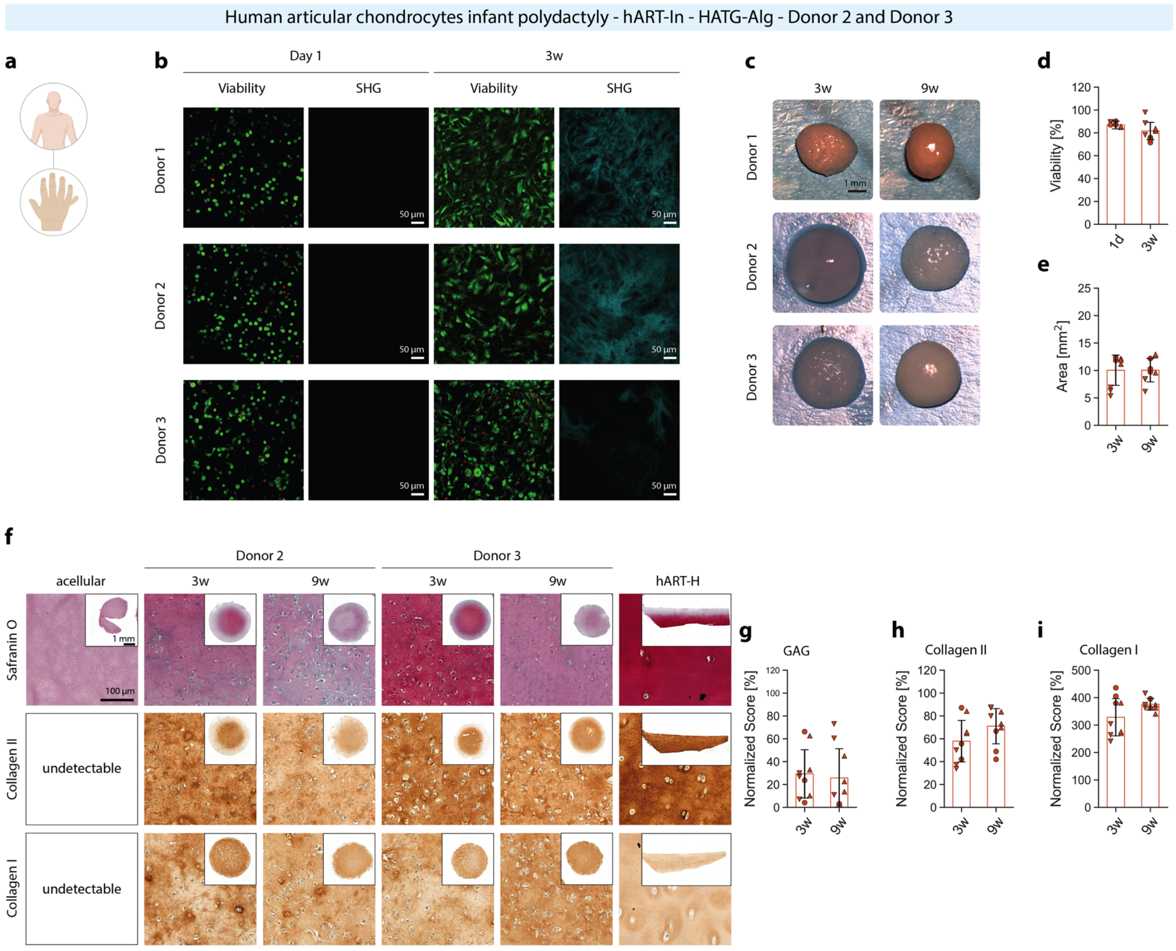


**Fig. S7 | hART-In in HATG-Alg.** (**a**) Schematic of hART-In sourced from polydactyly digits (age: 1.3 ± 0.6 years, age range: 1-2 years, sex: 3 males). (**b**) Viability and second harmonic generation of hART-In grafts at day 1 and 3 weeks and (**d**) viability quantification. (**c**) Gross appearance of hART-In grafts after 3 and 9 weeks and (**e**) graft area. (**f**) Histological analysis of acellular HATG-Alg, hART-In grafts matured after 3 and 9 weeks, and healthy human articular cartilage (hART-H). (**g**) GAG, (**h**) collagen II and (**i**) collagen I histological score breakdown between 3- and 9-weeks samples. Scale bar in viability and second harmonic generation images: 50 µm; scale bar in gross pictures of the samples and in histological overview images: 1 mm; scale bar in high-magnification images: 100 µm. hART-In: $n_d$ = 3, $n_s$ = 3.

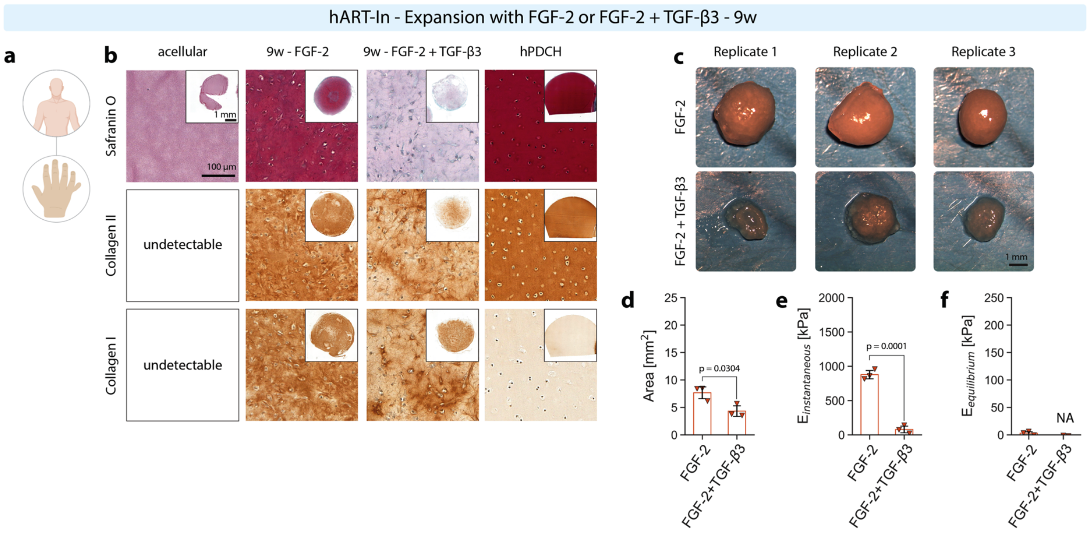


**Fig. S8 | hART-In expanded with FGF-2 and expanded with FGF-2+TGF-β3 in HATG-Alg.** (**a**) Schematic of hART-In sourced from polydactyly digits (age: 1.3 ± 0.6 years, age range: 1-2 years, sex: 3 males). (**b**) Histological analysis of hART-In grafts matured for 9 weeks after expansion with FGF-2 10 ng $mL^{-1}$ or FGF-2+TGF-β3 (both 10 ng $mL^{-1}$), compared to acellular HATG-Alg and polydactyly human cartilage (hART-In). (**c**) Gross appearance of hART-In grafts after expansion with FGF-2 or FGF-2+TGF-β3 and (**d**) graft area. (**e**) Instantaneous and (**f**) equilibrium moduli of hART-In grafts after expansion with FGF-2 or FGF-2+TGF-β3. Scale bar in gross pictures of the samples and in histological overview images: 1 mm; scale bar in high-magnification images: 100 μm. $n_d$ = 1, $n_s$ = 3.

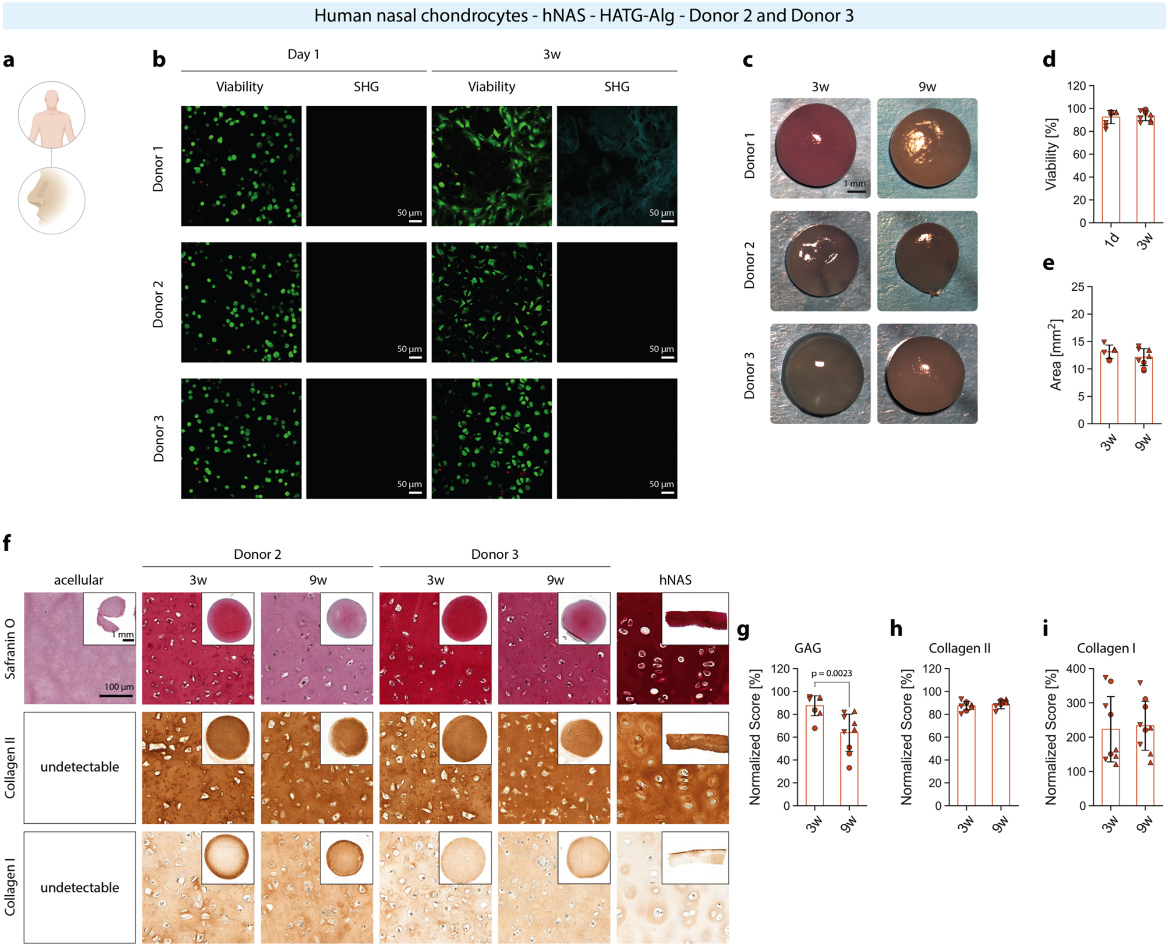


**Fig. S9 | hNAS in HATG-Alg.** (**a**) Schematic of hNAS sourced from nasal reconstructive surgery (age: 24.7 ± 6.1 years, age range: 18-30 years). (**b**) Viability and second harmonic generation of hNAS grafts at day 1 and 3 weeks and (**d**) viability quantification. (**c**) Gross appearance of hNAS grafts after 3 and 9 weeks and (**e**) graft area. (**f**) Histological analysis of acellular HATG-Alg, hNAS grafts matured after 3 and 9 weeks, and human nasal cartilage (hNAS). (**g**) GAG, (**h**) collagen II and (**i**) collagen I histological score breakdown between 3- and 9-weeks samples. Scale bar in viability and second harmonic generation images: 50 µm; scale bar in gross pictures of the samples and in histological overview images: 1 mm; scale bar in high-magnification images: 100 µm. hNAS: $n_d$ = 3, $n_s$ = 3.

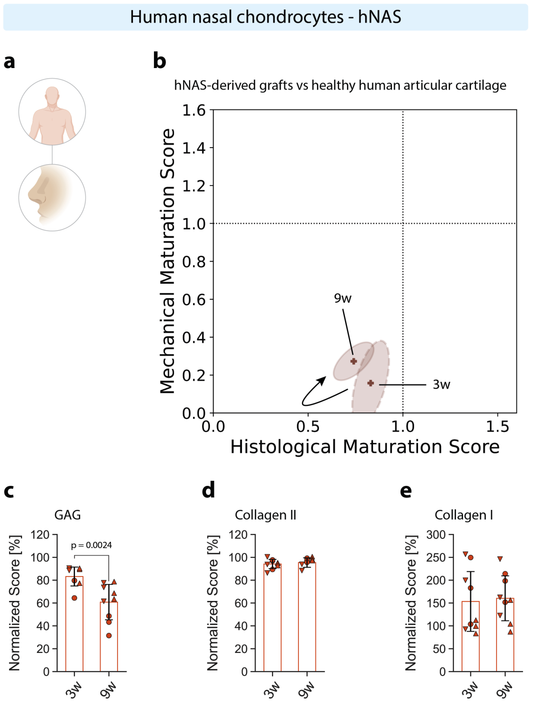


**Fig. S10 | hNAS – Maturation trajectory vs native healthy human articular cartilage.** (**a**) Schematic of hNAS sourced from nasal reconstructive surgery (age: 24.7 ± 6.1 years, age range: 18-30 years). (**b**) Scoring trajectory of hNAS maturation from 3 to 9 weeks compared to healthy human articular cartilage. (**c**) GAG, (**d**) collagen II and (**e**) collagen I histological score breakdown between 3- and 9-weeks hNAS samples normalized to healthy human articular cartilage.

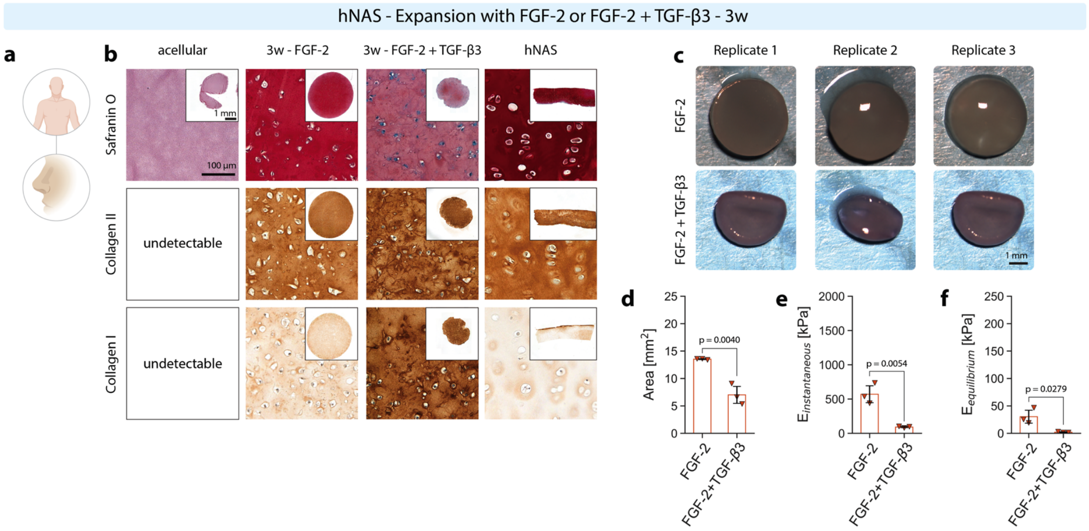


**Fig. S11 | hNAS expanded with FGF-2 and expanded with FGF-2+TGF-β3 in HATG-Alg.** (**a**) Schematic of hNAS sourced from nasal reconstructive surgery (age: 24.7 ± 6.1 years, age range: 18-30 years). (**b**) Histological analysis of hNAS grafts matured for 9 weeks after expansion with FGF-2 10 ng $mL^{-1}$ or FGF-2+TGF-β3 (both 10 ng $mL^{-1}$), compared to acellular HATG-Alg and human nasal cartilage (hNAS). (**c**) Gross appearance of hNAS grafts after expansion with FGF-2 or FGF-2+TGF-β3 and (**d**) graft area. (**e**) Instantaneous and (**f**) equilibrium moduli of hNAS grafts after expansion with FGF-2 or FGF-2+TGF-β3. Scale bar in gross pictures of the samples and in histological overview images: 1 mm; scale bar in high-magnification images: 100 µm. $n_d = 1$, $n_s = 3$.

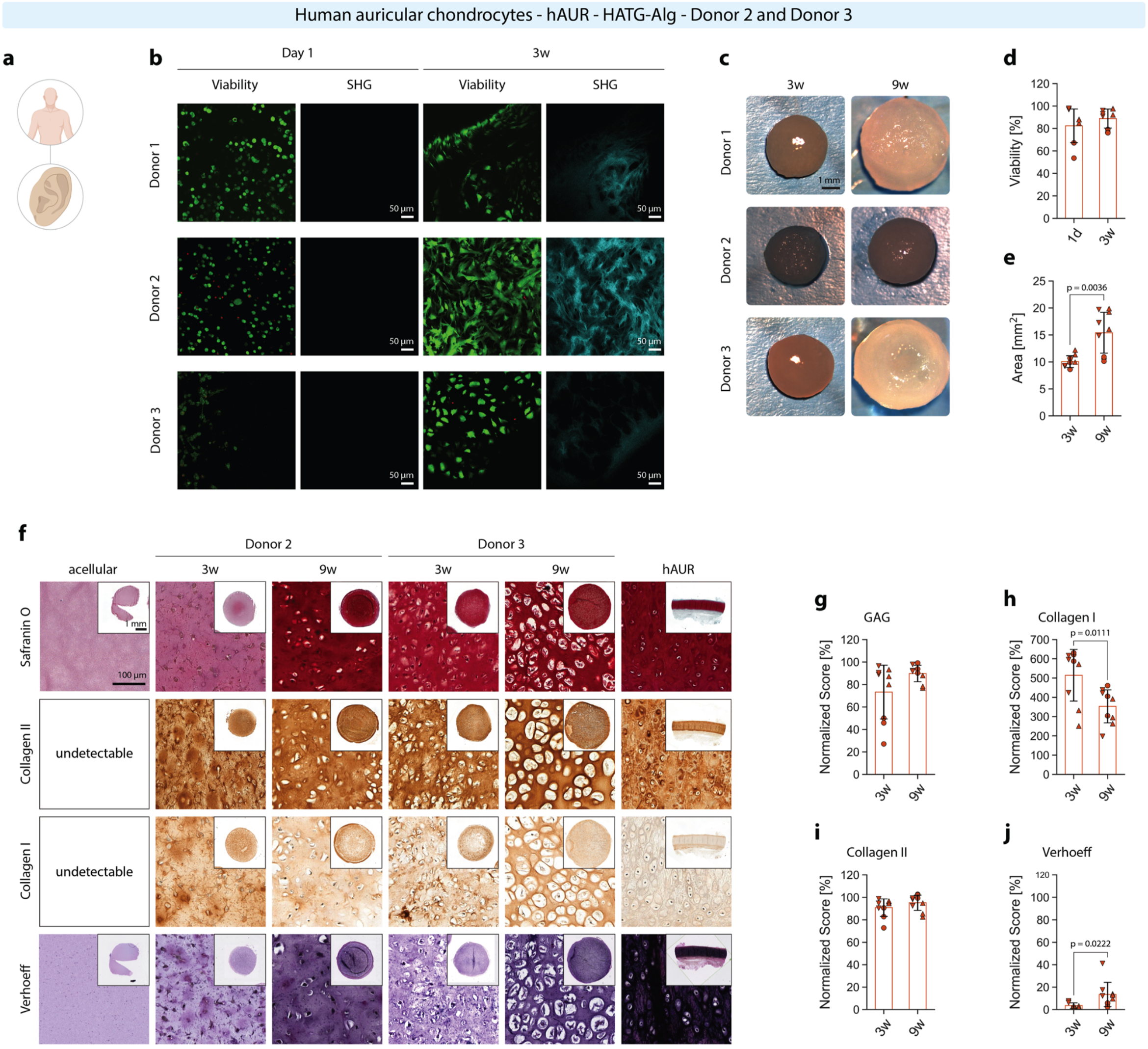


**Fig. S12 | hAUR in HATG-Alg.** (**a**) Schematic of hAUR sourced from otoplasty surgery (age: 11.3 ± 4.7 years, age range: 6-15 years, sex: 2 females, 1 male). (**b**) Viability and second harmonic generation of hAUR grafts at day 1 and 3 weeks and (**d**) viability quantification. (**c**) Gross appearance of hAUR grafts after 3 and 9 weeks and (**e**) graft area. (**f**) Histological analysis of acellular HATG-Alg, hAUR grafts matured after 3 and 9 weeks, and human auricular cartilage (hAUR). (**g**) GAG, (**h**) collagen I, (**i**) collagen II and (**j**) elastin histological score breakdown between 3- and 9-weeks samples. Scale bar in viability and second harmonic generation images: 50 µm; scale bar in gross pictures of the samples and in histological overview images: 1 mm; scale bar in high-magnification images: 100 µm. hAUR: $n_d = 3$, $n_s = 3$.

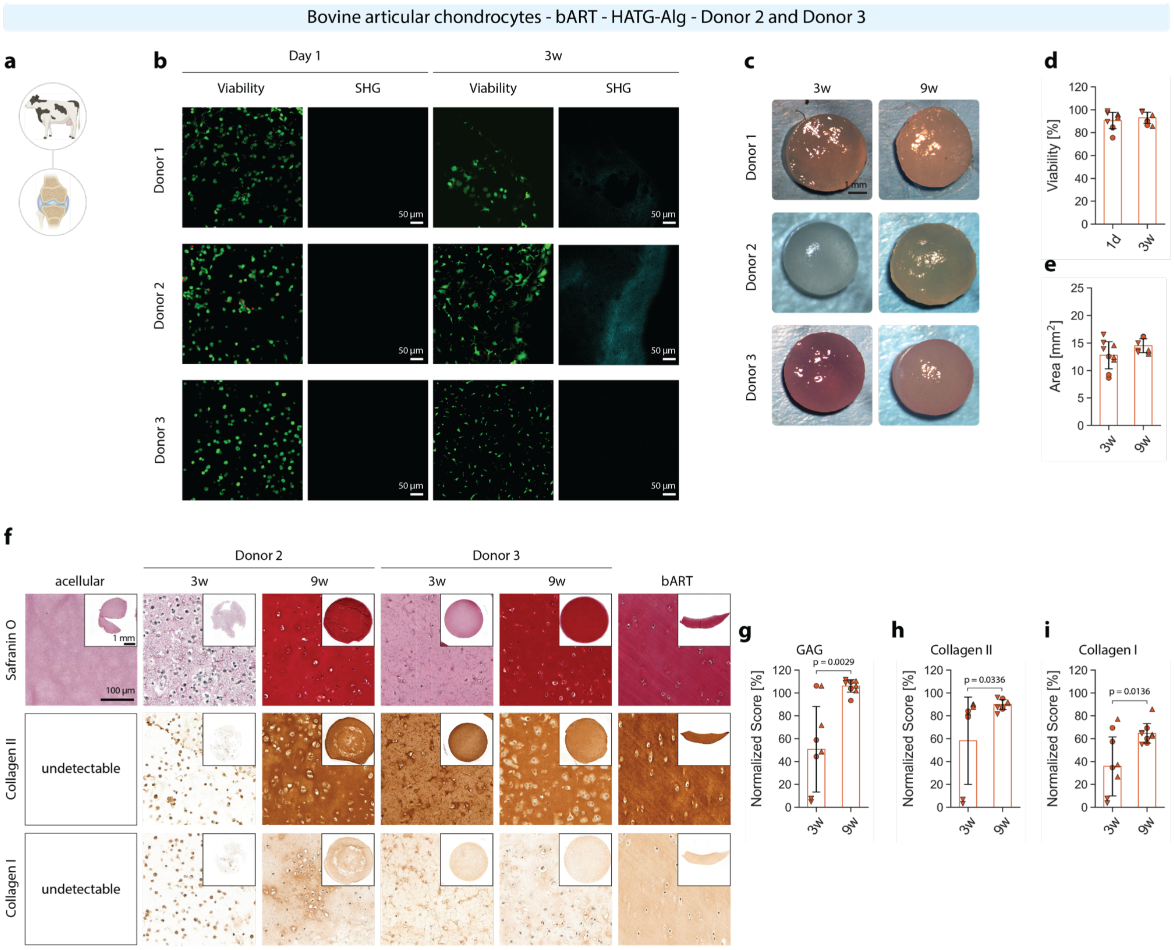


**Fig. S13 | bART in HATG-Alg.** (**a**) Schematic of bART sourced from bovine knee joints (~ 6-month-old calves). (**b**) Viability and second harmonic generation of bART grafts at day 1 and 3 weeks and (**d**) viability quantification. (**c**) Gross appearance of bART grafts after 3 and 9 weeks and (**e**) graft area. (**f**) Histological analysis of acellular HATG-Alg, bART grafts matured after 3 and 9 weeks, and bovine articular cartilage (bART). (**g**) GAG, (**h**) collagen II and (**i**) collagen I histological score breakdown between 3- and 9-weeks samples. Scale bar in viability and second harmonic generation images: 50 µm; scale bar in gross pictures of the samples and in histological overview images: 1 mm; scale bar in high-magnification images: 100 µm. bART: $n_d$ = 3, $n_s$ = 3.

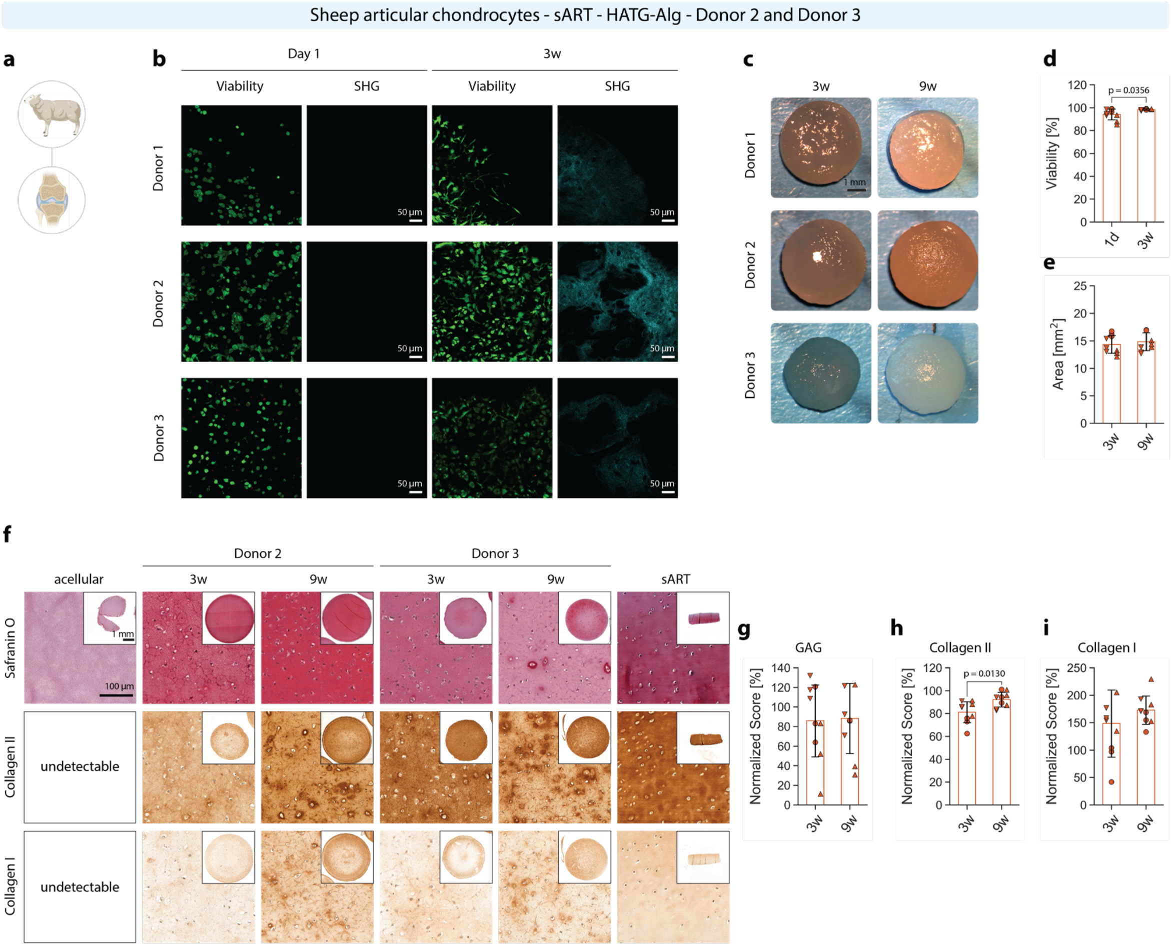


**Fig. S14 | sART in HATG-Alg.** (**a**) Schematic of sART sourced from sheep knee joints (~ 3-year-old sheep). (**b**) Viability and second harmonic generation of sART grafts at day 1 and 3 weeks and (**d**) viability quantification. (**c**) Gross appearance of sART grafts after 3 and 9 weeks and (**e**) graft area. (**f**) Histological analysis of acellular HATG-Alg, sART grafts matured after 3 and 9 weeks, and sheep articular cartilage (sART). (**g**) GAG, (**h**) collagen II and (**i**) collagen I histological score breakdown between 3- and 9-weeks samples. Scale bar in viability and second harmonic generation images: 50 µm; scale bar in gross pictures of the samples and in histological overview images: 1 mm; scale bar in high-magnification images: 100 µm. sART: $n_d$ = 3, $n_s$ = 3.

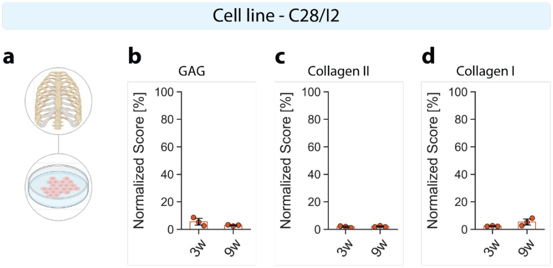


**Fig. S15 | C28/I2 cell line – Histology scoring.** (**a**) Schematic of the C28/I2 cell line, derived from costal cartilage chondrocytes which were subsequently immortalized. (**b**) GAG, (**c**) collagen II and (**d**) collagen I histological score breakdown between 3- and 9-weeks samples. $n_d$ = 1, $n_s$ = 3.

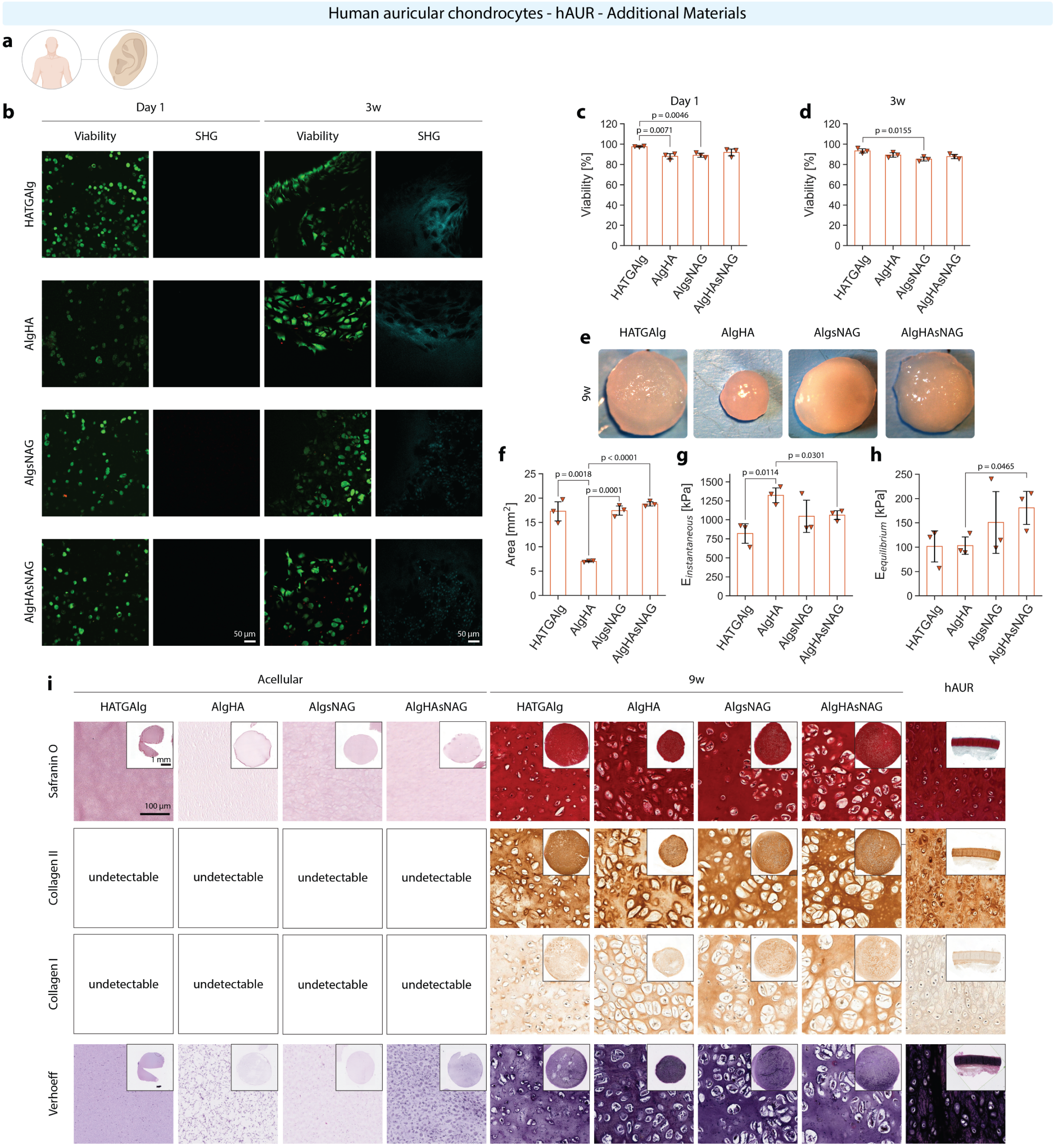


**Fig. S16 | hAUR in alginate-based materials.** (**a**) Schematic of hAUR sourced from otoplasty surgery (age: 11.3 ± 4.7 years, age range: 6-15 years, sex: 2 females, 1 male). (**b**) Viability and second harmonic generation of hAUR grafts in HATG-Alg, Alg-HA, Alg-sNAG and Alg-HA-sNAG at day 1 and 3 weeks and (**c**, **d**) viability quantification. (**e**) Gross appearance of hAUR grafts after 3 and 9 weeks in HATG-Alg, Alg-HA, Alg-sNAG and Alg-HA-sNAG and (**f**) graft area. (**g**) Instantaneous and (**h**) equilibrium modulus of hAUR grafts encapsulated in HATG-Alg, Alg-HA, Alg-sNAG and Alg-HA-sNAG. (**i**) Histological analysis of acellular HATG-Alg, Alg-HA, Alg-sNAG and Alg-HA-sNAG controls, hAUR grafts matured in all 4 alginate-based materials after 3 and 9 weeks, and human auricular cartilage (hAUR). $n_d = 1$, $n_s = 3$.

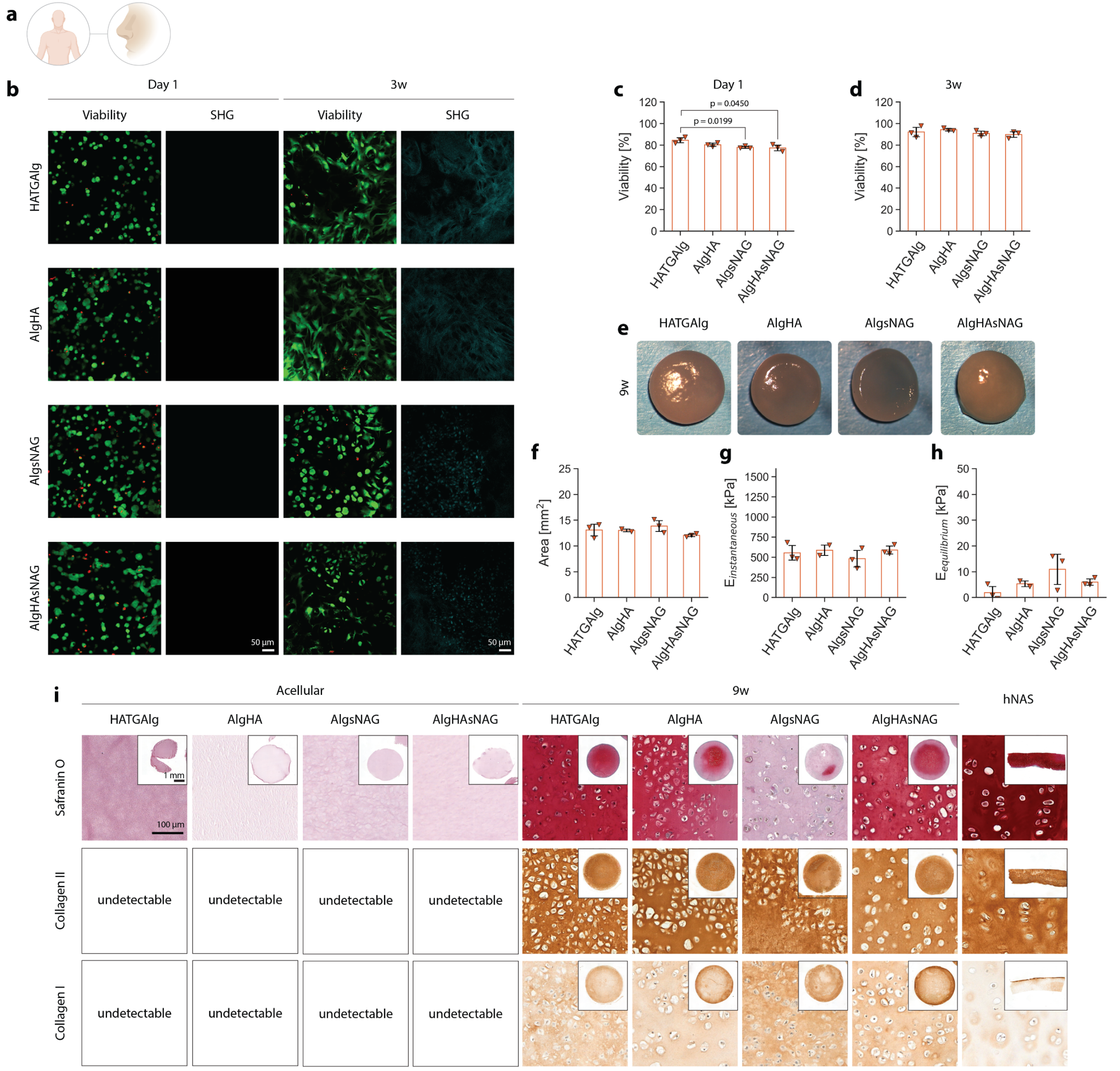


**Fig. S17 | hNAS in alginate-based materials.** (**a**) Schematic of hNAS sourced from nasal reconstructive surgery (age: 24.7 ± 6.1 years, age range: 18-30 years). (**b**) Viability and second harmonic generation of hNAS grafts in HATG-Alg, Alg-HA, Alg-sNAG and Alg-HA-sNAG at day 1 and 3 weeks and (**c**, **d**) viability quantification. (**e**) Gross appearance of hNAS grafts after 3 and 9 weeks in HATG-Alg, Alg-HA, Alg-sNAG and Alg-HA-sNAG and (**f**) graft area. (**g**) Instantaneous and (**h**) equilibrium modulus of hNAS grafts encapsulated in HATG-Alg, Alg-HA, Alg-sNAG and Alg-HA-sNAG. (**i**) Histological analysis of acellular HATG-Alg, Alg-HA, Alg-sNAG and Alg-HA-sNAG controls, hNAS grafts matured in all 4 alginate-based materials after 3 and 9 weeks, and human nasal cartilage (hNAS). $n_d = 1$, $n_s = 3$.

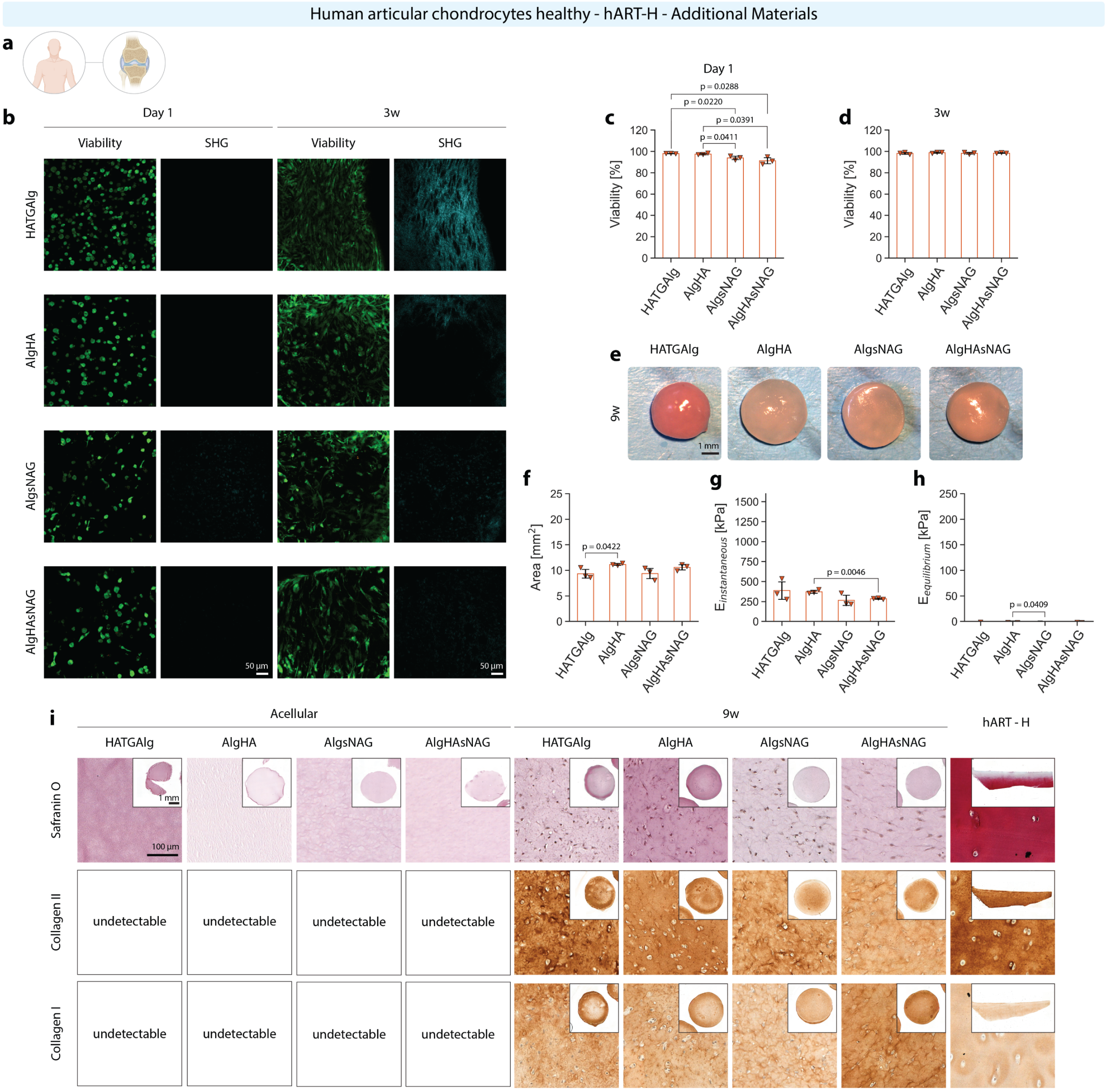


**Fig. S18 | hART-H in alginate-based materials.** (**a**) Schematic of hART-H sourced from healthy knee joints (age: 58.0 ± 6.6 years, age range: 51-64 years, sex: 1 female, 2 males). (**b**) Viability and second harmonic generation of hART-H grafts in HATG-Alg, Alg-HA, Alg-sNAG and Alg-HA-sNAG at day 1 and 3 weeks and (**c**, **d**) viability quantification. (**e**) Gross appearance of hART-H grafts after 3 and 9 weeks in HATG-Alg, Alg-HA, Alg-sNAG and Alg-HA-sNAG and (**f**) graft area. (**g**) Instantaneous and (**h**) equilibrium modulus of hART-H grafts encapsulated in HATG-Alg, Alg-HA, Alg-sNAG and Alg-HA-sNAG. (**i**) Histological analysis of acellular HATG-Alg, Alg-HA, Alg-sNAG and Alg-HA-sNAG controls, hART-H grafts matured in all 4 alginate-based materials after 3 and 9 weeks, and healthy human articular cartilage (hART-H). $n_d$ = 1, $n_s$ = 3.

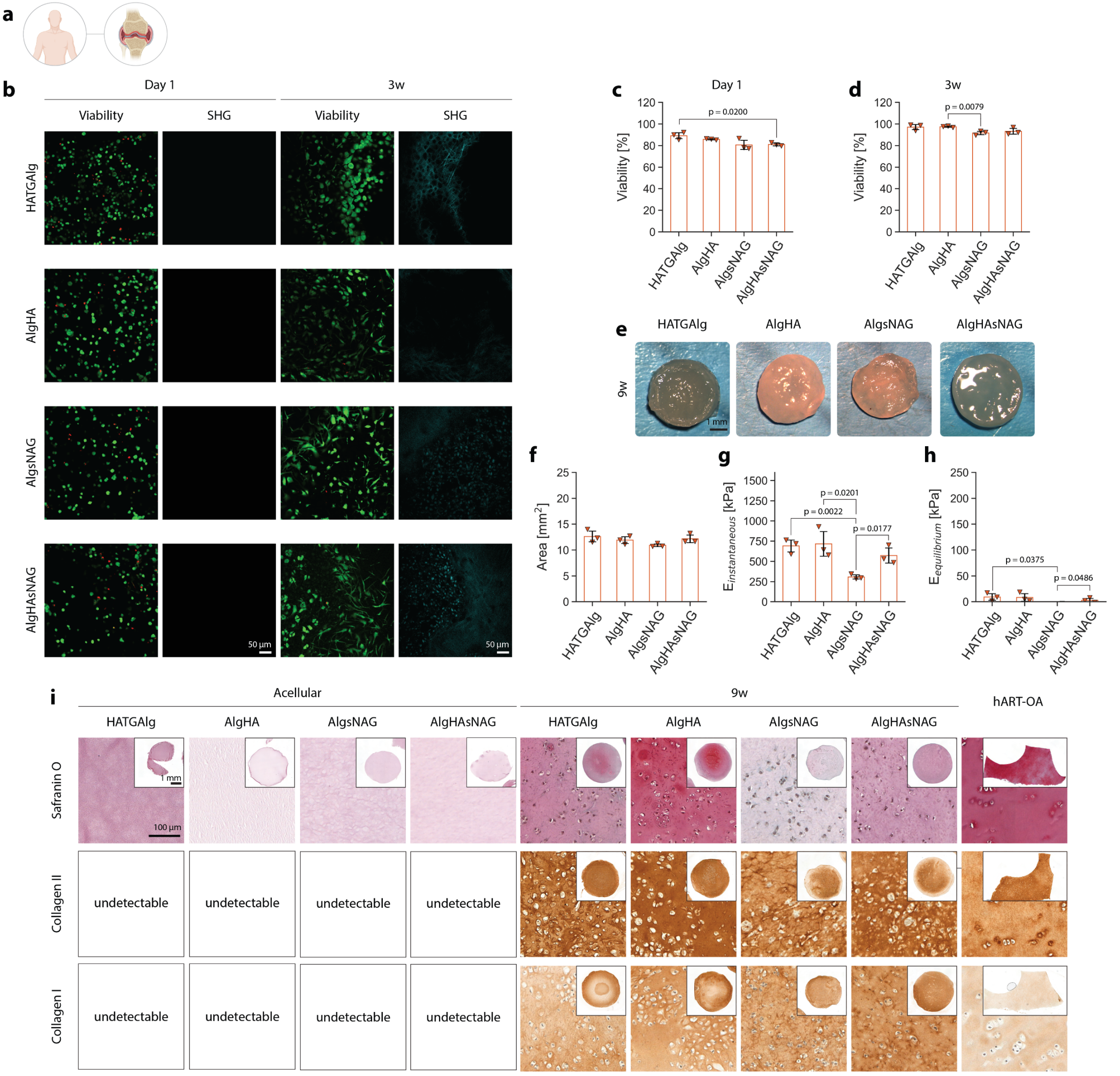


**Fig. S19 | hART-OA in alginate-based materials.** (**a**) Schematic of hART-OA sourced from osteoarthritic knee joints (age: 49.0 ± 19.2 years, age range: 27-62 years, sex: 2 females, 1 male). (**b**) Viability and second harmonic generation of hART-OA grafts in HATG-Alg, Alg-HA, Alg-sNAG and Alg-HA-sNAG at day 1 and 3 weeks and (**c**, **d**) viability quantification. (**e**) Gross appearance of hART-OA grafts after 3 and 9 weeks in HATG-Alg, Alg-HA, Alg-sNAG and Alg-HA-sNAG and (**f**) graft area. (**g**) Instantaneous and (**h**) equilibrium modulus of hART-OA grafts encapsulated in HATG-Alg, Alg-HA, Alg-sNAG and Alg-HA-sNAG. (**i**) Histological analysis of acellular HATG-Alg, Alg-HA, Alg-sNAG and Alg-HA-sNAG controls, hART-OA grafts matured in all 4 alginate-based materials after 3 and 9 weeks, and OA human articular cartilage (hART-OA). $n_d$ = 1, $n_s$ = 3.

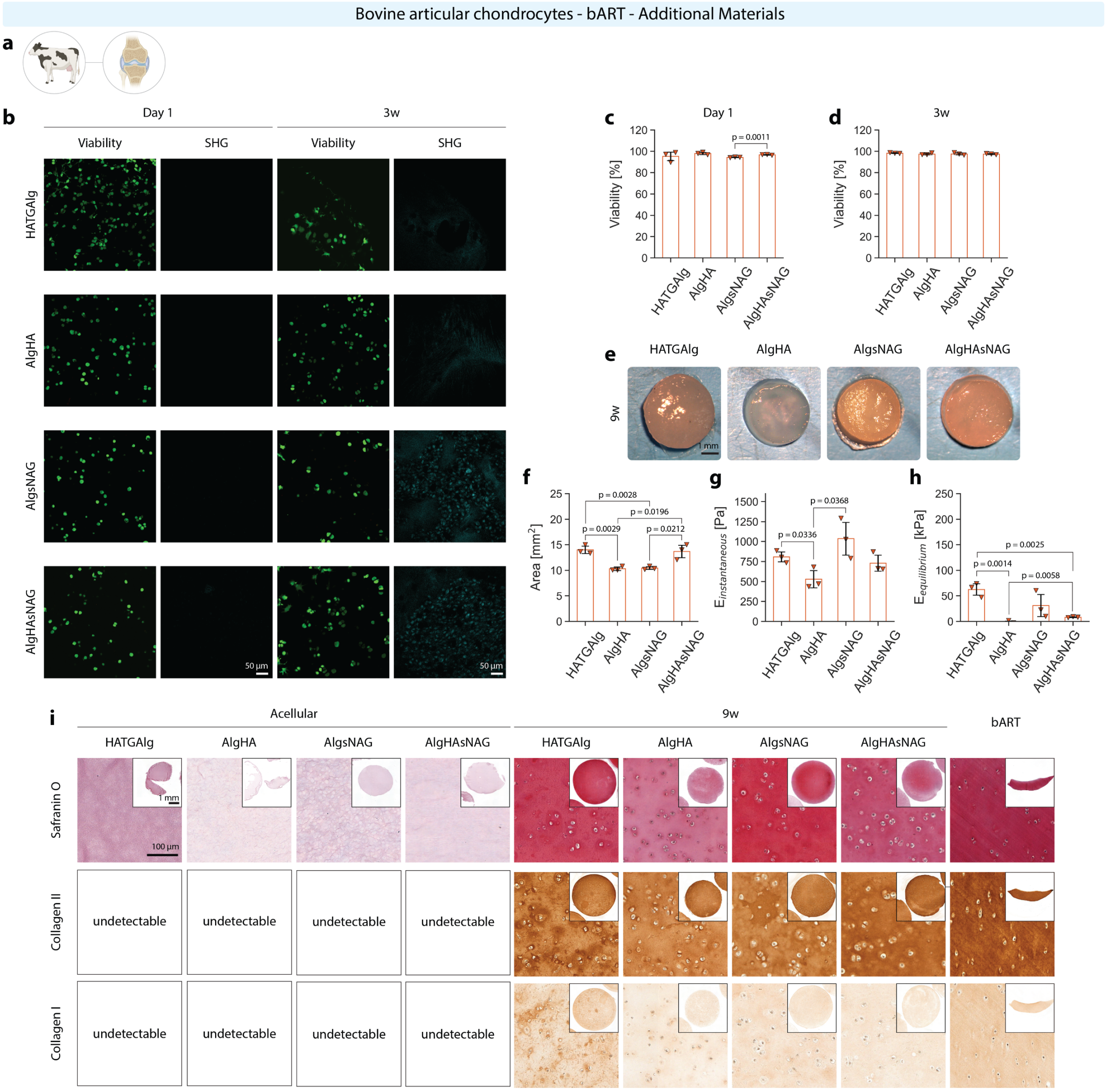


**Fig. S20 | bART in alginate-based materials.** (**a**) Schematic of bART sourced from bovine knee joints (~ 6-month-old calves). (**b**) Viability and second harmonic generation of bART grafts in HATG-Alg, Alg-HA, Alg-sNAG and Alg-HA-sNAG at day 1 and 3 weeks and (**c**, **d**) viability quantification. (**e**) Gross appearance of bART grafts after 3 and 9 weeks in HATG-Alg, Alg-HA, Alg-sNAG and Alg-HA-sNAG and (**f**) graft area. (**g**) Instantaneous and (**h**) equilibrium modulus of bART grafts encapsulated in HATG-Alg, Alg-HA, Alg-sNAG and Alg-HA-sNAG. (**i**) Histological analysis of acellular HATG-Alg, Alg-HA, Alg-sNAG and Alg-HA-sNAG controls, bART grafts matured in all 4 alginate-based materials after 3 and 9 weeks, and bovine articular cartilage (bART). $n_d$ = 1, $n_s$ = 3.

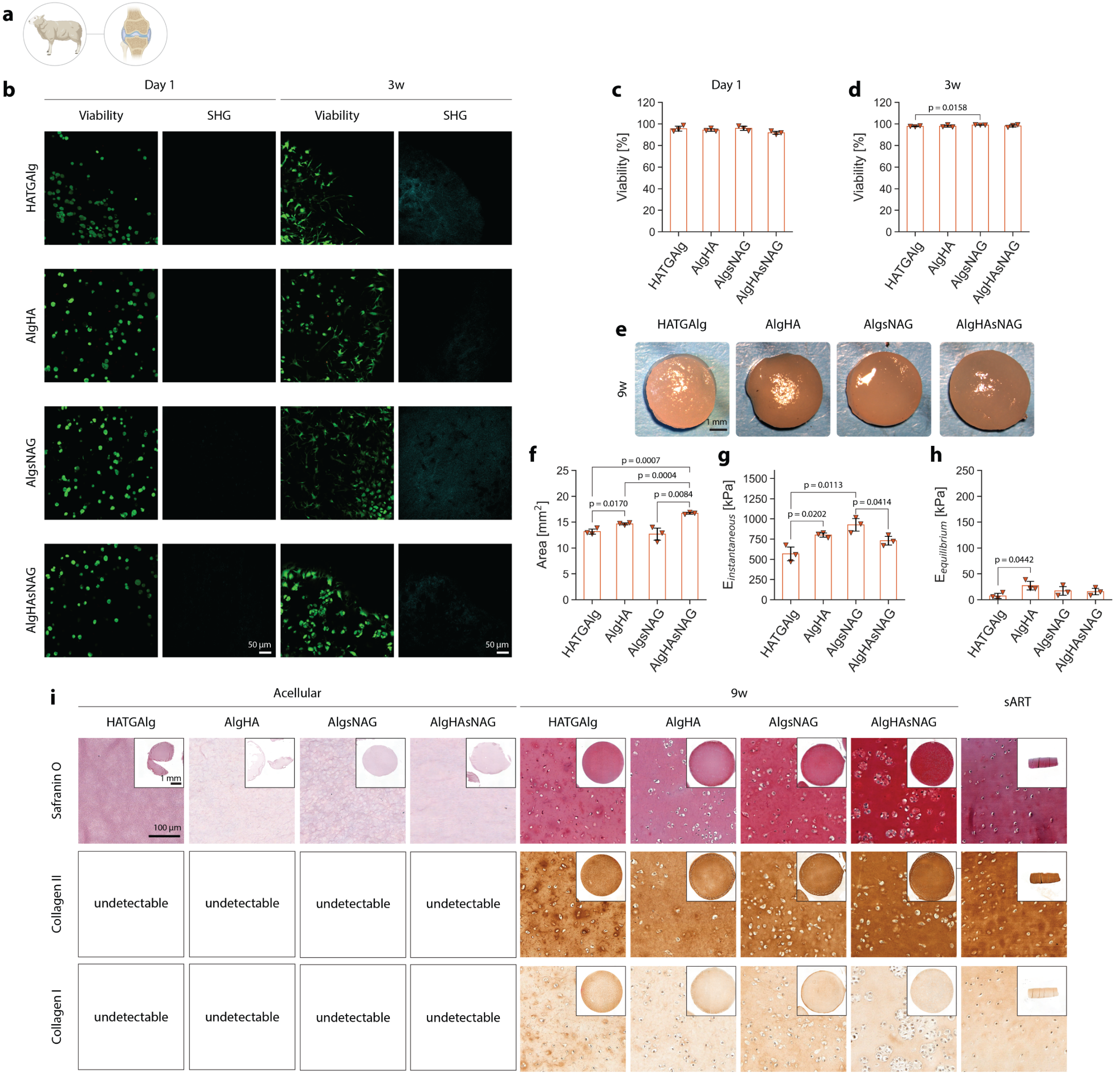


**Fig. S21 | sART in alginate-based materials.** (**a**) Schematic of sART sourced from sheep knee joints (~ 3-year-old sheep). (**b**) Viability and second harmonic generation of sART grafts in HATG-Alg, Alg-HA, Alg-sNAG and Alg-HA-sNAG at day 1 and 3 weeks and (**c**, **d**) viability quantification. (**e**) Gross appearance of sART grafts after 3 and 9 weeks in HATG-Alg, Alg-HA, Alg-sNAG and Alg-HA-sNAG and (**f**) graft area. (**g**) Instantaneous and (**h**) equilibrium modulus of sART grafts encapsulated in HATG-Alg, Alg-HA, Alg-sNAG and Alg-HA-sNAG. (**i**) Histological analysis of acellular HATG-Alg, Alg-HA, Alg-sNAG and Alg-HA-sNAG controls, sART grafts matured in all 4 alginate-based materials after 3 and 9 weeks, and sheep articular cartilage (sART). $n_d$ = 1, $n_s$ = 3.
